# PD-linked LRRK2 G2019S mutation impairs astrocyte morphology and synapse maintenance via ERM hyperphosphorylation

**DOI:** 10.1101/2023.04.09.536178

**Authors:** Shiyi Wang, Ryan Baumert, Gabrielle Séjourné, Dhanesh Sivadasan Bindu, Kylie Dimond, Kristina Sakers, Leslie Vazquez, Jessica L. Moore, Christabel Xin Tan, Tetsuya Takano, Maria Pia Rodriguez, Nick Brose, Luke Bradley, Reed Lessing, Scott H. Soderling, Albert R. La Spada, Cagla Eroglu

## Abstract

Astrocytes are highly complex cells that mediate critical roles in synapse formation and maintenance by establishing thousands of direct contacts with synapses through their perisynaptic processes. Here, we found that the most common Parkinsonism gene mutation, LRRK2 G2019S, enhances the phosphorylation of the ERM proteins (Ezrin, Radixin, and Moesin), components of the perisynaptic astrocyte processes in a subset of cortical astrocytes. The ERM hyperphosphorylation was accompanied by decreased astrocyte morphological complexity and reduced excitatory synapse density and function. Dampening ERM phosphorylation levels in LRRK2 G2019S mouse astrocytes restored both their morphology and the excitatory synapse density in the anterior cingulate cortex. To determine how LRRK2 mutation impacts Ezrin interactome, we used an *in vivo* BioID proteomic approach, and we found that astrocytic Ezrin interacts with Atg7, a master regulator of autophagy. The Ezrin/Atg7 interaction is inhibited by Ezrin phosphorylation, thus diminished in LRRK2 G2019S astrocytes. Importantly, the Atg7 function is required to maintain proper astrocyte morphology. Our data provide a molecular pathway through which the LRRK2 G2019S mutation alters astrocyte morphology and synaptic density in a brain-region-specific manner.

## INTRODUCTION

Most synapses in the central nervous system are in direct contact with perisynaptic astrocyte processes (PAPs)[1], forming the “tripartite synapse”[1]. At the tripartite synapse, astrocytes clear excess neurotransmitters, maintain ion homeostasis, and secrete factors regulating synapse development and function[2]. To accomplish these crucial functions, astrocytes develop elaborate morphologies to tile the entire brain parenchyma[3–5]. Astrocyte morphogenesis and synaptogenesis are interdependent and bidirectional[6, 7]. In neurological disorders like Parkinson’s Disease (PD), astrocytes become reactive, undergoing morphological and molecular changes[8, 9]. Recent studies suggested that astrocyte dysfunction contributes to PD[10–12]. However, the underlying mechanisms are largely unknown.

Mutations in the leucine-rich repeat kinase 2 (*LRRK2*) gene cause familial PD[13, 14], with LRRK2 G2019S being the most common[15, 16]. It is largely accepted that LRRK2 G2019S is a gain-of-function mutation that increases LRRK2’s kinase activity[17–20]. Brain RNA-seq databases suggested that LRRK2 expressed in astrocytes[21, 22]. Previous studies also suggested ERM proteins (Ezrin, Radixin and Moesin) as LRRK2 substrates[18, 23, 24]. ERM proteins are structural components of perisynaptic astrocyte processes (PAPs), with Ezrin being the most enriched in expression[25, 26]. Previous studies suggested that LRRK2 phosphorylates a conserved threonine residue present in all ERM proteins[18, 23, 24]. When phosphorylated, ERM proteins change conformation and link the F-actin cytoskeleton to the cell membrane[27, 28]. This switch is thought to play an important role in organizing cell morphology[28, 29]. Among the ERM proteins, Ezrin is specifically and highly enriched in astrocytes and is known to localize at the PAPs preferentially[25]. Genetic ablation or downregulation of Ezrin has been shown to alter astrocyte morphology *in vivo*[30–33].

Here, we tested whether LRRK2 regulates astrocyte morphology and synaptic densities using the LRRK2 G2019S knockin (^ki/ki^) mouse model[34, 35] in conjunction with *in vivo* astrocyte-specific genetic, proteomic, electrophysiological, and biochemical approaches. We found that, in the prefrontal cortex of humans and mice carrying the LRRK2 G2019S mutation, a subset of astrocytes has increased ERM (Ezrin, Radixin, and Moesin) phosphorylation levels. This phospho-ERM change accompanies three observations: Astrocytes have reduced morphological complexity; The anterior cingulate cortex (ACC) has decreased excitatory (VGLUT1/PSD95) synapse density; The primary motor cortex (MOp) has increased inhibitory (VGAT/Gephyrin) synapse density. By overexpression of an Ezrin mutant that cannot be phosphorylated in astrocytes, we dampened astrocytic ERM hyperphosphorylation. This reduction in astrocytic ERM phosphorylation restored both LRRK2 G2019S astrocyte morphology and the ACC excitatory synapse density. However, this phospho-ERM manipulation did not rescue the MOp inhibitory synapse changes. We further compared *in vivo* Ezrin interactomes in WT and LRRK2 G2019S^ki/ki^ mouse cortical astrocytes. We discovered that Ezrin interacts with a master regulator of autophagy, Atg7. This interaction is weakened by Ezrin’s phosphorylation. We also found that Atg7 is required for astrocyte morphological complexity. Knocking down Atg7 in LRRK2 G2019S astrocytes partially restores their morphology. This study shows that LRRK2-ERM pathways are important for astrocytes to reach their morphological complexity. This study also hinted that astrocyte morphological changes correlate with synapse density changes in a brain region-specific manner.

## RESULTS

### ERM phosphorylation is increased in the prefrontal cortices of mice and humans carrying the LRRK2 G2019S mutation

To study how the LRRK2 G2019S mutation impacts astrocytes in the prefrontal cortex, a cortical region linked to PD, we stained sections from the frontal cortices of age and sex-matched control and LRRK2 G2019S mutation-carrying PD patients (Supplemental Table 1) with glial fibrillary acidic protein (GFAP, a marker for astrocyte branches) and phosphorylated ERM (Phospho-ERM, which are enriched in PAPs) (Figure 1A). Remarkably, the astrocytic organization, determined by GFAP staining, was severely disrupted, and GFAP levels were significantly elevated (∼42%) in PD patients carrying the LRRK2 G2019S mutation, indicative of reactive gliosis (Figure 1B-C). Unexpectedly, the phospho-ERM signal was increased by 7-fold in the frontal cortices of PD patients carrying the LRRK2 G2019S mutation (Figure 1B-C). To determine if this ERM hyperphosphorylation is cell type-specific we co-stained for phospho-ERM with astrocyte marker S100β and neuronal dendrite marker MAP2. We found a significant increase in phospho-ERM signal colocalized with both S100β and MAP2 (Figure S1A-C).

**Figure 1:**
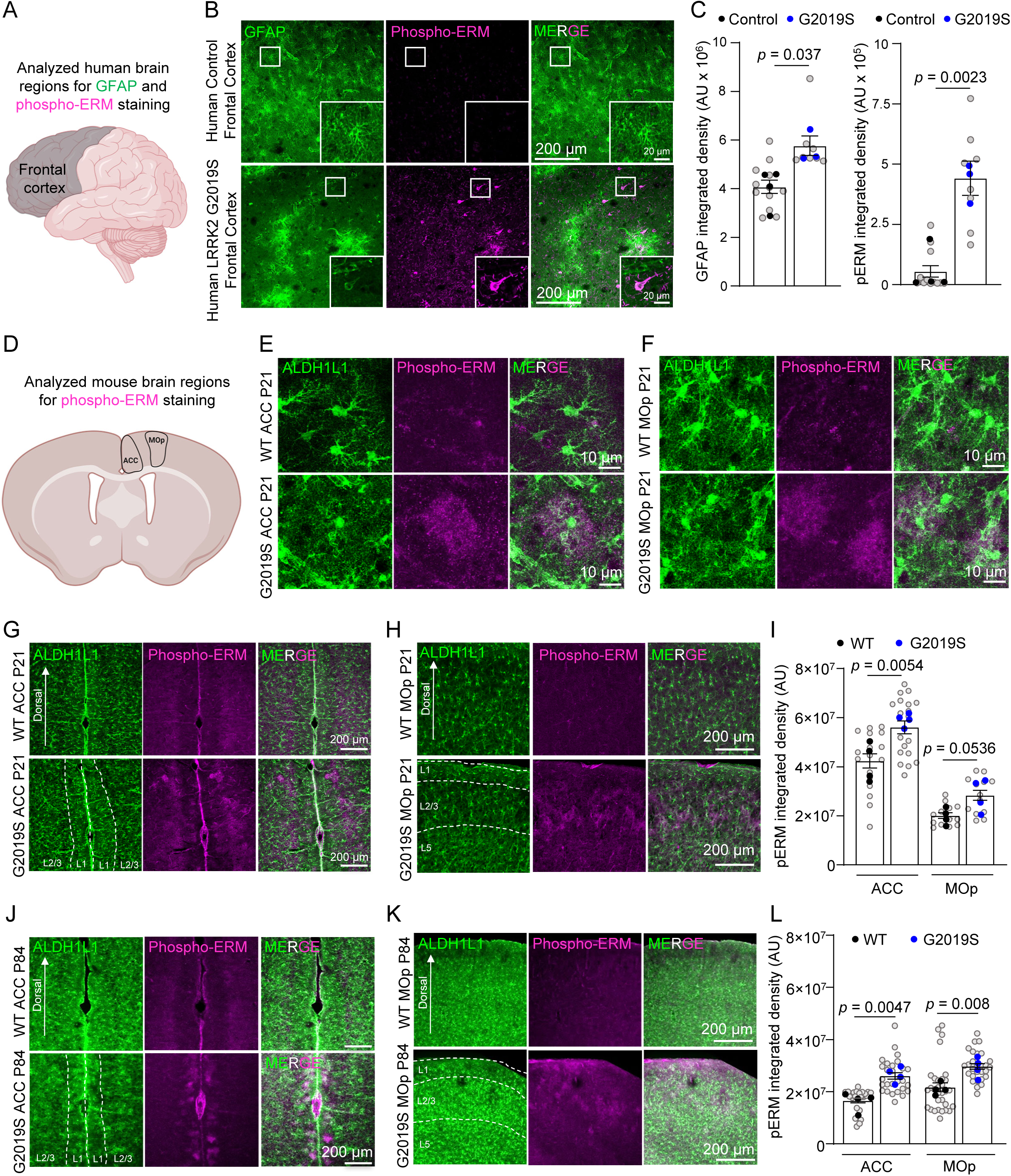
ERM phosphorylation is impaired in PD patients carrying LRRK2 G2019S mutation. (A) Schematic of human frontal cortex regions analyzed for phospho-ERM staining. (B) Representative confocal images of GFAP (green) and phospho-ERM (purple) in the frontal cortex of human controls (n = 4; 3M, 1F) and LRRK2 G2019S mutation carriers (n = 3; 2M, 1F), aged >80 years. Scale bar, 200 μm. (C) Quantification of GFAP integrated density in (B). GFAP: t (5) = 2.822, *p* = 0.037; phospho-ERM: t (5) = 5.695, *p* = 0.0023. Grey dots: individual images; black dots: control averages; blue dots: mutation carrier averages. (D) Schematic of mouse anterior cingulate cortex (ACC) and primary motor cortex (MOp) analyzed for phospho-ERM staining. (E-F) Representative confocal images of phospho-ERM in the ACC and the MOp of WT or LRRK2 G2019S^ki/ki^ Aldh1L1-eGFP mice at P21. Scale bar, 10 μm. (G-H) Representative confocal images of phospho-ERM in the ACC and MOp of WT (n = 4; 2M, 2F) or LRRK2 G2019S^ki/ki^ Aldh1L1-eGFP mice (n = 4; 2M, 2F) at P21. Scale bar, 200 μm. (I) Quantification of phospho-ERM integrated density in (G-H). For the ACC, nested t-test, unpaired two-tailed t-test. t (6) = 4.247, *p* = 0.0054. For the MOp, nested t-test, unpaired two-tailed t-test. t (6) = 2.392, *p* = 0.0536. (J-K) Representative confocal images of phospho-ERM (purple) in the ACC and MOp of WT (n = 4; 2M, 2F) or LRRK2 G2019S^ki/ki^ Aldh1L1-eGFP (n = 4; 2M, 2F) mice at P84. Scale bar, 200 μm. (L) Quantification of phospho-ERM integrated density in (J-K), For the ACC, nested t-test, unpaired Two-tailed t-test. t (6) = 4.383, *p* = 0.0047. For the MOp, nested t-test, unpaired Two-tailed t-test. t (6) = 3.898, *p* = 0.0080. Grey dots: individual images; black dots: WT averages; blue dots: mutant averages.

To determine whether ERM hyperphosphorylation is an early event in LRRK2 G2019S PD pathology, we used LRRK2 G2019S^ki/ki^ mice, in which the human G2019S mutation was introduced into exon 41 of the LRRK2 gene[34, 35]. To determine if ERM hyperphosphorylation localizes to astrocytes, we crossed these mice to an astrocyte reporter line carrying the Aldh1L1-eGFP transgene, which labels all astrocytes by GFP[36]. We examined the phospho-ERM levels in coronal brain sections from 21-day-old (P21, juvenile) and 12-week-old (P84, adult) WT or LRRK2 G2019S^ki/ki^ mice. Compared to WT mice, LRRK2 G2019S^ki/ki^ mice across both age groups have a brain region-specific and heterogeneous phospho-ERM increase (Figure 1D-L). This phospho-ERM increase was localized to layers (L)1 and L2-3 of the anterior cingulate cortex (ACC) and the primary motor cortex (MOp) (Figure 1E-L; Figure S2A-C) but was absent in both the primary sensory cortex (SSp) and the dorsal medial striatum (DMS) (Figure S2D-I). In P21 LRRK2 G2019S^ki/ki^ mice, the phospho-ERM integrated densities in the ACC and MOp L1 and L2-3 were increased by ∼32% (ACC) and ∼41% (MOp) compared to the WTs (Figure 1I); In P84 LRRK2 G2019S^ki/ki^ mice, the phospho-ERM integrated densities were increased by ∼57% (ACC) and ∼37% (MOp) compared to the WTs (Figure 1L). In the ACC and MOp L1 and L2/3 of LRRK2 G2019S^ki/ki^ mice, phospho-ERM was significantly increased and colocalized with S100β and MAP2 but not Iba1 (microglia marker), with a predominant localization to astrocytes in the ACC and MOp of LRRK2 G2019S^ki/ki^ mice (Figure S3A-C). We also confirmed the specificity of the phospho-ERM antibody by treating brain sections from both mice and human samples with lambda protein phosphatase (λ-PPase). λ-PPase dephosphorylates proteins[37]. In both cases, the phospho-ERM signal was eliminated after λ-PPase treatment, confirming the phospho-ERM antibody specificity (Figure S4A-F).

In the human samples, phospho-ERM elevation was observed concurrently with reactive gliosis (elevated GFAP staining). To determine if, in LRRK2 G2019S^ki/ki^ mice, there were reactive astrocytes detectable at early ages, we stained sections from the same P21 mouse brains with an anti-GFAP antibody. In the mouse cortex, grey matter astrocytes do not display GFAP staining unless there is reactive gliosis[38, 39]. We did not detect any increase in the levels of GFAP (Figure S5A-C). These data indicate that, in LRRK2 G2019S^ki/ki^ mice, phospho-ERM upregulation does not occur concurrently with reactive astrogliosis. Moreover, distinct from the P21 and P84 LRRK2 G2019S^ki/ki^ mice where phospho-ERM was restricted to a subset of astrocytes in a brain region-specific manner, in the human patient brains (∼80 years old), phospho-ERM accumulation was present equally in both astrocytes and neurons. These data suggest that ERM hyperphosphorylation is an early and astrocyte-specific event in LRRK2 G2019S^ki/ki^ mouse cortices; however, is more widespread in human patient brains.

### LRRK2 G2019S mutation alters excitatory and inhibitory synapse densities and functions in the cortex

Previous studies did not observe dopaminergic neuron loss or motor dysfunction in the LRRK2 G2019S^ki/ki^ mice[40, 41]; however, synapse dysfunction at cortico-striatal circuits was reported in adult mice[42]. Despite the critical role of LRRK2 in cortico-striatal synapses and PD-related cortical circuits[43], whether LRRK2 G2019S affects cortical synapse densities and function has not been explored. Understanding these cortical synaptic abnormalities is crucial, as they may underlie the early cognitive deficits in PD. Next, we investigated if LRRK2 G2019S mutation alters synaptic densities in the ACC and MOp L1 and L2-3 (Figure 2A), regions in which ERM hyperphosphorylation within astrocytes was observed.

**Figure 2:**
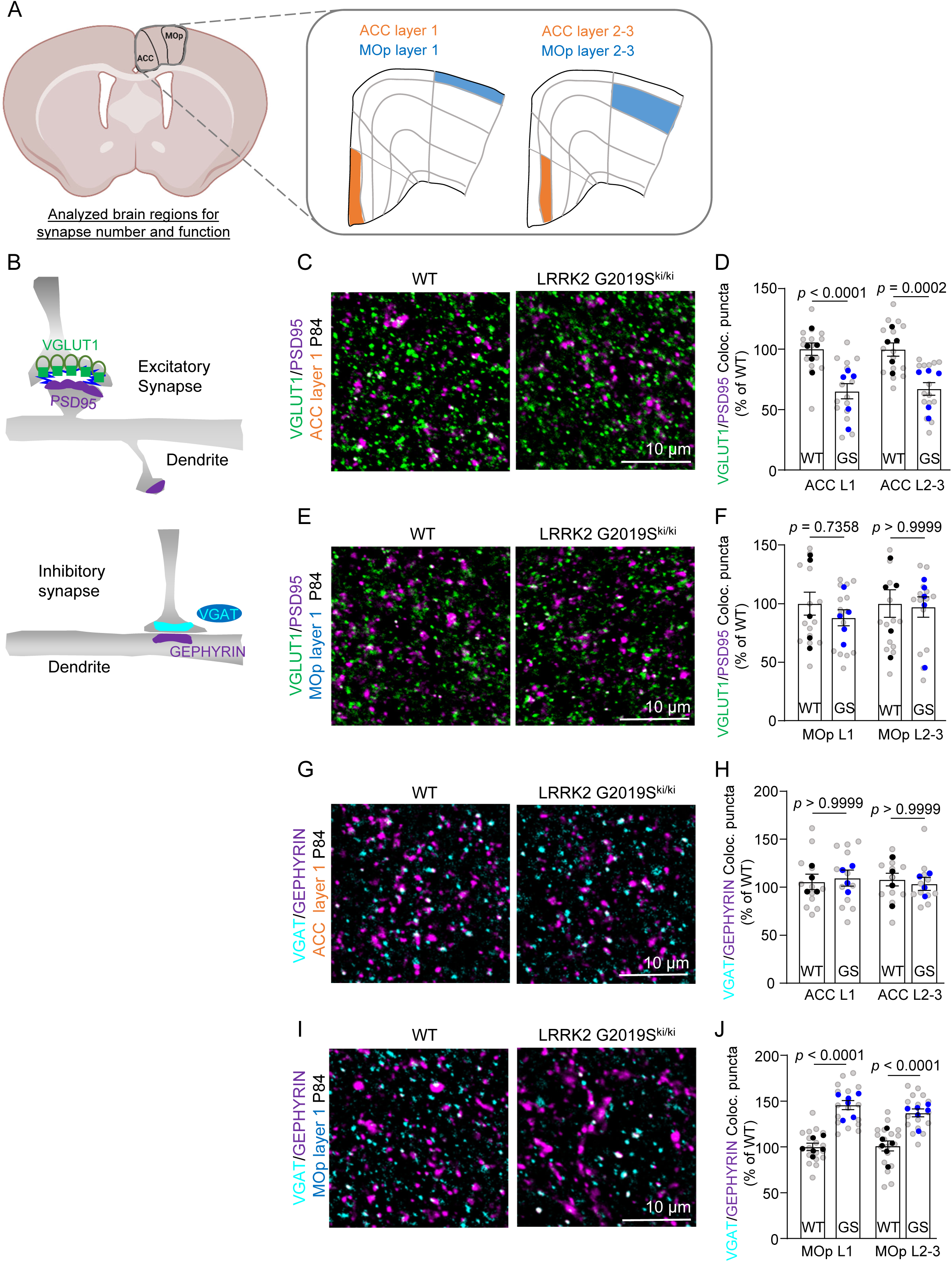
LRRK2 G2019S affects excitatory and inhibitory synapse densities in the ACC and MOp. (A-B) Schematics of analyzed brain regions (ACC and MOp) and methods for quantifying VGluT1-PSD95 and VGAT-GEPHYRIN colocalized puncta. (C) Representative images of VGluT1-PSD95 staining in ventral ACC L1 of WT and LRRK2 G2019S^ki/ki^ mice at P84. Scale bar, 10 μm. (D) Quantification of VGluT1-PSD95 puncta in ventral ACC L1 and L2-3, normalized to WT means. n = 5/group (3M, 2F). Nested One-way ANOVA: [F(3, 56) = 12.48, *p* < 0.0001]. Bonferroni’s test: L1 (*p* < 0.0001), L2-3 (*p* = 0.0002). (E) Representative images of VGluT1-PSD95 staining in MOp L1 at P84. Scale bar, 10 μm. (F) Quantification of VGluT1-PSD95 puncta in MOp L1 and L2-3. n = 5/group (3M, 2F). No significant differences (Bonferroni’s test: *p* > 0.05). (G) Representative images of VGAT-GEPHYRIN staining in ventral ACC L1. Scale bar, 10 μm. (H) Quantification of VGAT-GEPHYRIN puncta in ACC L1 and L2-3, normalized to WT means. n = 4/group (2M, 2F). No significant differences (Bonferroni’s test: *p* > 0.05). (I) Representative images of VGAT-GEPHYRIN staining in MOp L1. Scale bar, 10 μm. (J) Quantification of VGAT-GEPHYRIN puncta in MOp L1 and L2-3. n = 6/group (3M, 3F). Nested One-way ANOVA: [F(3, 68) = 25.88, *p* < 0.0001]. Bonferroni’s test: L1 (*p* < 0.0001), L2-3 (*p* < 0.0001). Grey dots: individual images; black dots: WT averages; blue dots: mutant averages.

We quantified the densities of excitatory and inhibitory intracortical synapses in WT and LRRK2 G2019S^ki/ki^ mice. To do so, we marked the excitatory synapses as the co-localization of the presynaptic Vesicular Glutamate Transporter 1 (VGlut1) and the Postsynaptic Density Protein (PSD95), whereas inhibitory synapses were labeled using presynaptic Vesicular GABA Transporter (VGAT), and postsynaptic gephyrin [44, 45] (Figure 2B). Structural synapses were identified as the apposition and co-localization of presynaptic and postsynaptic markers, which are in two distinct neuronal compartments (i.e., axons and dendrites) and would only appear to co-localize at synapses due to their proximity[46].

Excitatory synapse density was quantified both at P21 and P84. In the ACC of the LRRK2 G2019S^ki/ki^ mice, there was a significant decrease in intracortical excitatory synapse density in both layers and ages we analyzed (∼30%) when compared to the WT controls (Figure 2C-D and Figure S6A-B). However, no differences in excitatory synapse density were observed between genotypes in the MOp (Figure 2E-F and Figure S6C-D). When we quantified the densities of inhibitory synapses in the MOp, we found significant differences between genotypes (∼40% increase). In contrast, in the ACC of WT and LRRK2 G2019S^ki/ki^ mice, no differences were observed for inhibitory synapse density (Figure 2G-J). These results revealed two changes in LRRK2 G2019S^ki/ki^ mouse brains: reduced excitatory synapse density in the ACC; increased inhibitory synapse density in the MOp. These data suggested a brain region-specific synapse density change associated with the LRRK2 G2019S mutation.

We then used electrophysiology to assess how altered synapse density in the ACC and MOp of LRRK2 G2019S^ki/ki^ mice affects synaptic function. We recorded miniature excitatory postsynaptic currents (mEPSCs) in the ACC L2-3 pyramidal neurons and miniature inhibitory postsynaptic currents (mIPSCs) in the MOp L2-3 pyramidal neurons from acute brain slices of P84 WT and LRRK2 G2019S^ki/ki^ mice. LRRK2 G2019S^ki/ki^ neurons displayed reduced mEPSC frequency by ∼65% and a corresponding right shift in the cumulative distributions of mEPSC inter-event intervals (Figure 3A-B) when compared to WT neurons, with no change in mEPSC amplitude (Figure 3C). mEPSC frequency changes correlate with synapse densities and/or the probability of presynaptic glutamate release[47], whereas mEPSC amplitude indicates the synaptic strength reflecting the number of AMPARs at the postsynapse[48]. Therefore, together with our anatomical data, these findings suggest that the LRRK2 G2019S mutation reduces the number of intracortical excitatory connections formed on L2-3 pyramidal neurons in the ACC without altering synaptic strength. In contrast, in the MOp, LRRK2 G2019S^ki/ki^ neurons exhibited a ∼72% increase in mIPSC frequency compared to WT, with no significant change in amplitude (Figure 3D-F). These data indicate that inhibitory synapse density is elevated in the MOp of the LRRK2 G2019S^ki/ki^ mice.

**Figure 3:**
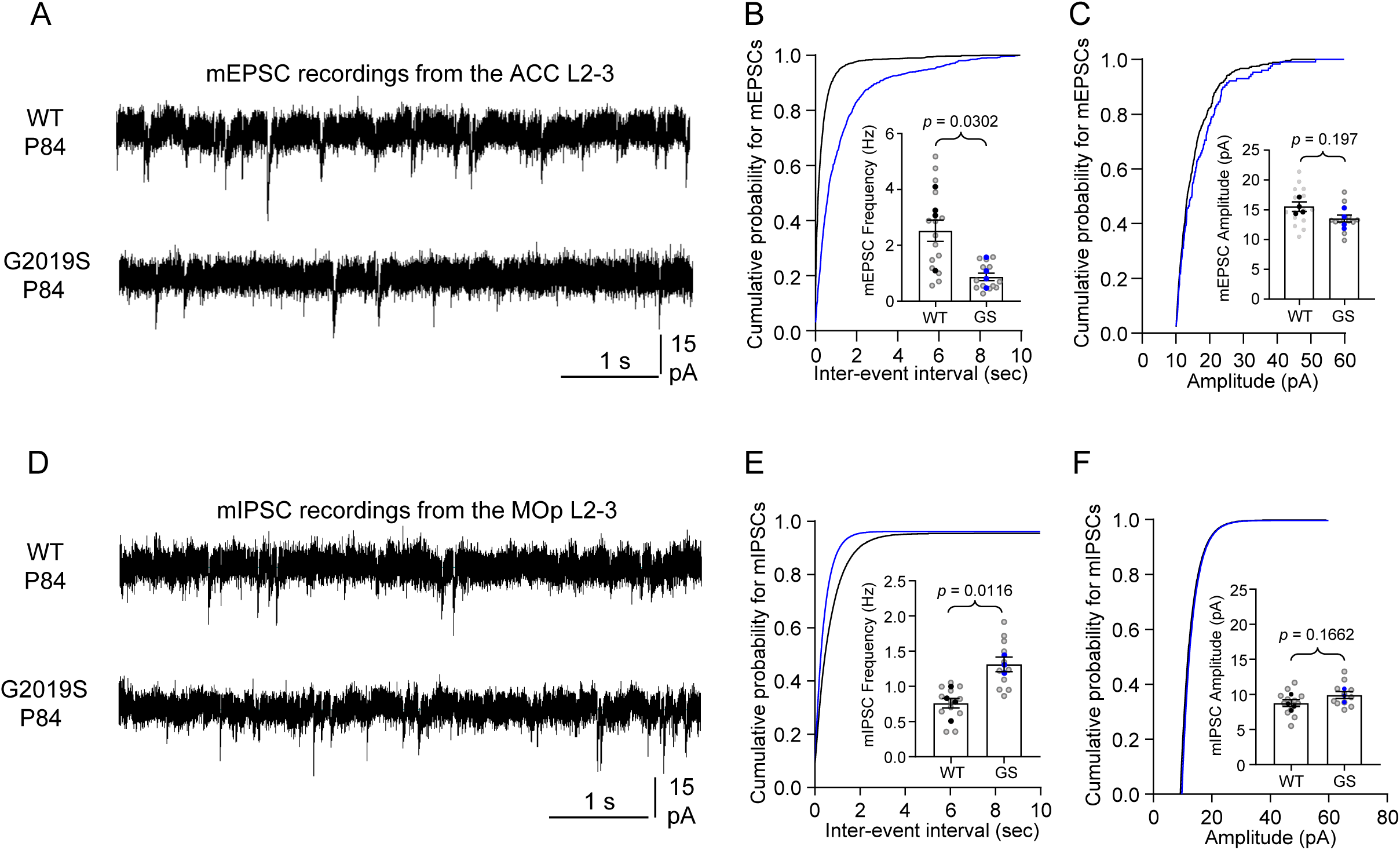
LRRK2 G2019S affects excitatory and inhibitory synapse function in the ACC and MOp. (A) Representative mEPSC traces from ventral ACC L2-3 pyramidal neurons in WT and LRRK2 G2019S^ki/ki^ mice. (B) Cumulative probability and quantification of mEPSC frequency: n = 15 (WT), 13 (LRRK2 G2019S^ki/ki^) neurons, 4 mice/genotype. Kolmogorov-Smirnov test: D = 0.504, *p* < 0.001. Mean frequency: WT (2.878 ± 0.6355), LRRK2 G2019S^ki/ki^ (0.9673 ± 0.2328). Unpaired t-test: t (6) = 2.823, *p* = 0.0302. (C) Cumulative probability and quantification of mEPSC amplitude: n = 14 (WT), 12 (LRRK2 G2019S^ki/ki^) neurons. Kolmogorov-Smirnov test: D = 0.194, *p* < 0.0001. Mean amplitude: WT (15.5 ± 0.6325), LRRK2 G2019S^ki/ki^ (13.9 ± 0.9017). Unpaired t-test: t (6) = 1.451, *p* = 0.197. (D) Representative mIPSC traces from MOp L2-3 pyramidal neurons in WT and LRRK2 G2019S^ki/ki^ mice. (E) Cumulative probability and quantification of mIPSC frequency: n = 12 (WT), 11 (LRRK2 G2019S^ki/ki^) neurons, 4 WT, 3 LRRK2 G2019S^ki/ki^ mice. Kolmogorov-Smirnov test: D = 0.424, *p* < 0.0001. Mean frequency: WT (0.7788 ± 0.1012), LRRK2 G2019S^ki/ki^ (1.301 ± 0.07291). Unpaired t-test: t (5) = 3.883, *p* = 0.0116. (F) Cumulative probability and quantification of mIPSC amplitude: n = 12 (WT), 11 (LRRK2 G2019S^ki/ki^) neurons. Kolmogorov-Smirnov test: D = 0.382, *p* < 0.0001. Mean amplitude: WT (8.633 ± 0.4883), LRRK2 G2019S^ki/ki^ (9.834 ± 0.5558). Unpaired t-test: t (5) = 1.62, *p* = 0.1662. Data are presented as mean ± s.e.m.

### Reducing ERM hyperphosphorylation in astrocytes restores excitatory synapse density and function in the ACC of adult LRRK2 G2019S^ki/ki^ mice

LRRK2 phosphorylates a conserved threonine residue present in all ERM proteins[18, 23]. We found that LRRK2 G2019S mutation increases ERM phosphorylation in astrocytes (Figure 1) and asked if dampening ERM phosphorylation in astrocytes could restore synapse densities in LRRK2 G2019S^ki/ki^ mice. To test this, we overexpressed a phospho-dead Ezrin mutant, which cannot be phosphorylated, in astrocytes via adenovirus (AAV) retro-orbital injection. We chose Ezrin because Ezrin is highly abundant and specifically enriched in astrocytes compared to its family members, Radixin and Moesin[21] (Figure S7A-B). Switching the amino acid threonine (T) at location 567 with alanine (A) results in a version of the protein that cannot be phosphorylated (Phospho-dead Ezrin, Figure S7C)[28, 29].

To determine if phospho-dead Ezrin can reduce ERM phosphorylation in astrocytes of LRRK2 G2019S^ki/ki^ mice, we retro-orbitally injected 9-week-old WT and LRRK2 G2019S^ki/ki^ mice with AAVs to express Hemagglutinin (HA)-tagged WT Ezrin or phospho-dead Ezrin under the control of the astrocyte-specific GfaABC1D promoter (Figure S7D). The GfaABC1D promoter ensures astrocyte-specific expression, as validated in prior work[49]. While AAV serotypes may infect multiple cell types, our data support astrocyte-specific expression based on promoter specificity and cell labeling. 3 weeks after injection, the mouse brains were harvested for histological analyses. The widespread viral expression of Ezrin in astrocytes was verified by staining with an anti-HA tag antibody (Figure 4A). While WT or phospho-dead Ezrin overexpression had no effect on ERM phosphorylation in WT brains, WT Ezrin increased phospho-ERM levels in LRRK2 G2019S^ki/ki^ astrocytes; phospho-dead Ezrin abolished this increase (Figure 4A-B). This indicates that the LRRK2 G2019S sensitizes astrocytes to LRRK2-mediated ERM phosphorylation. To determine the specificity of phospho-dead Ezrin, we examined the hyperphosphorylation of another LRRK2 substrate, Rab10 (Thr 73)[19, 50],[51]. We found higher Rab10 phosphorylation in LRRK2 G2019S^ki/ki^ brain sections than in WT mice. However, neither WT nor phospho-dead Ezrin overexpression impacted Rab10 phosphorylation (Figure S8), showing that phospho-dead Ezrin selectively reduces ERM hyperphosphorylation in astrocytes.

**Figure 4:**
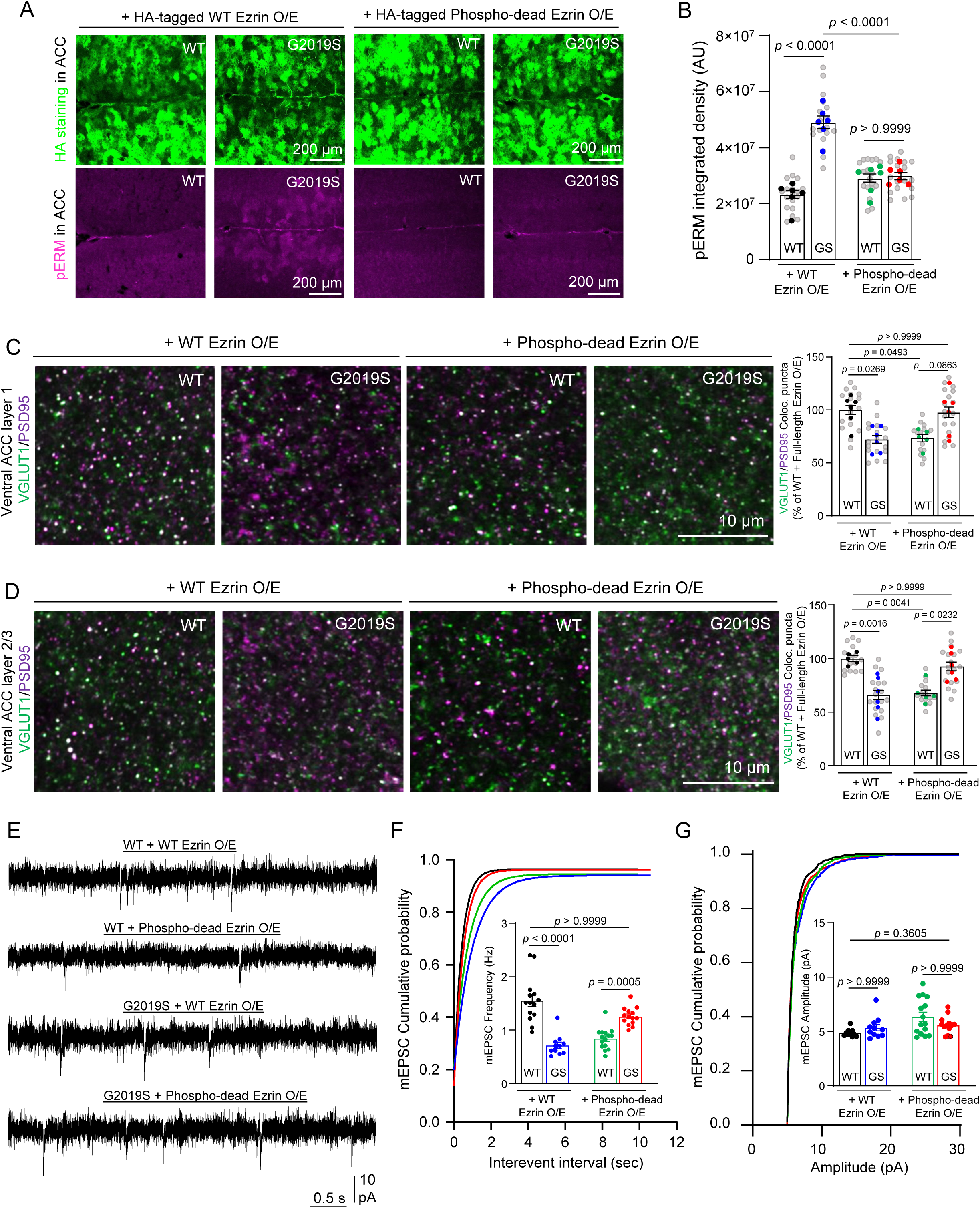
Overexpression of phospho-dead Ezrin in adult LRRK2 G2019S^ki/ki^ astrocytes restores excitatory synapse number and function in the ventral ACC. (A) Representative ventral ACC images of WT and LRRK2 G2019S^ki/ki^ mice injected with AAV-HA-WT or Phospho-dead EZRIN, stained for HA and phospho-ERM (P84). Scale bar, 200 μm. (B) Quantification of phospho-ERM intensity (n = 6/group, 3 males, 3 females). One-way ANOVA [F(3, 20) = 31.62, *p* < 0.0001], Bonferroni’s test: significant differences between WT + WT EZRIN vs. LRRK2 + WT EZRIN (*p* < 0.0001), and LRRK2 + WT EZRIN vs. LRRK2 + Phospho-dead EZRIN (*p* < 0.0001). No significant differences for other comparisons (*p* > 0.05). (C-D) Representative ACC L1 and L2-3 images of VGluT1/PSD95 staining (P84). Scale bar, 10 μm. Quantification of co-localized puncta normalized to WT + WT EZRIN (n = 5–6/group). L1: One-way ANOVA [F(3, 19) = 5.882, *p* = 0.0051], Bonferroni’s test: significant differences among WT + WT EZRIN, LRRK2 + WT EZRIN, and their Phospho-dead counterparts (*p* < 0.05). L2-3: One-way ANOVA [F(3, 18) = 10.45, *p* = 0.0003], Bonferroni’s test: significant group differences (*p* < 0.05). (E) Representative mEPSC traces from L2-3 pyramidal neurons. (F) Synaptic event frequency quantification (n = 11–15 neurons from 4 mice/group). Kruskal-Wallis test [H(3) = 36.83, *p* < 0.0001], Dunn’s posthoc: significant differences between WT + WT EZRIN vs. all other groups (*p* < 0.0001), except LRRK2 + Phospho-dead EZRIN (*p* > 0.9999). (G) Synaptic event amplitude (n = 11–15 neurons from 4 mice/group). Kruskal-Wallis test [H(3) = 7.051, *p* = 0.0703]; no significant differences across groups (*p* > 0.05). Data are mean ± s.e.m. Data are mean ± s.e.m.

We next tested whether reducing ERM hyperphosphorylation in astrocytes could rescue synaptic deficits in LRRK2 G2019S^ki/ki^ ACC and MOp. We found that, in WT ACC, WT Ezrin overexpression in astrocytes had no effect on excitatory synapse density (Figure S9). However, compared to WT mice overexpressing WT Ezrin, overexpression of phospho-dead Ezrin in WT astrocytes reduced excitatory synapse density by ∼30% (Figure 4C-D). In contrast, in LRRK2 G2019S^ki/ki^ astrocytes, overexpression of phospho-dead Ezrin increased the LRRK2 G2019S^ki/ki^ ACC excitatory synapse density. This increase brings the excitatory synapse density of LRRK2 G2019S^ki/ki^ ACC similar to those of WT ACC overexpressing astrocytic WT Ezrin (Figure 4C-D). We also found that, in the MOp L1 and L2-3 of LRRK2 G2019S^ki/ki^ mice, neither the WT nor the phospho-dead Ezrin overexpression in adult astrocytes restored inhibitory synapse density back to the levels in the WT MOp overexpressed with WT Ezrin (Figure S10). These data indicate that phospho-dead Ezrin expression in adult WT or LRRK2 G2019S^ki/ki^ astrocytes is sufficient to modify excitatory synapse density in the ACC but cannot alter inhibitory synapse density in the MOp. It is plausible that LRRK2 G2019S mutation impairs inhibitory synapses in the MOp by altering the number or connectivity of cortical interneurons through a developmental mechanism. Thus, manipulation in adults is insufficient to rectify this inhibitory synapse phenotype.

Because astrocytic overexpression of phospho-dead Ezrin impacted excitatory synapse density in the ACC of WT and LRRK2 G2019S^ki/ki^ mice, we next performed whole-cell patch-clamp recordings from ACC L2-3 pyramidal neurons transduced with the same AAVs. In LRRK2 G2019S^ki/ki^ ACC L2-3, overexpression of WT Ezrin in astrocytes did not rescue the decreased frequency of mEPSCs. However, overexpression of phospho-dead Ezrin in LRRK2 G2019S^ki/ki^ astrocytes restored the frequency up to the levels of WT mice expressing WT Ezrin (Figure 4E-F). Interestingly, in WT ACC L2-3, overexpression of phospho-dead Ezrin in WT astrocytes significantly reduced the frequency of mEPSCs (∼54%) (Figure 4E-F). Posthoc comparisons confirmed no significant differences in mean mEPSC amplitudes across all 4 conditions (Figure 4G).

Altogether, these electrophysiological and neuroanatomical analyses reveal that the LRRK2 G2019S mutation dysregulates proper excitatory and inhibitory synapse densities and functions in the cortex in a brain region-specific manner. These results also pinpoint a link between ERM hyperphosphorylation in LRRK2 G2019S astrocytes (Figure 1) and the excitatory synaptic density change in the ACC (Figures 2 and 3). Our results suggest that balancing ERM phosphorylation in astrocytes plays a role in maintaining excitatory synapse density and function in the ACC.

### LRRK2 regulates astrocyte morphological complexity *in vivo* through ERM phosphorylation

In the brain, astrocytes have numerous functions[2]. To accomplish these functions, astrocytes develop elaborate morphologies[3–5]. Astrocyte morphology is a crucial indicator for its function. We next asked whether ERM hyperphosphorylation in LRRK2 G2019S astrocytes would impair their morphological complexity. To answer this question, we sparsely labeled astrocytes with a membrane-targeted fluorescent protein (mCherry-CAAX) using postnatal astrocyte labeling by electroporation (PALE). To image and reconstruct the morphology of individual whole astrocytes, we collected 100-µm thick mouse brain sections (Figure S11A). We focused our analyses of astrocyte territory size and complexity on the ACC and MOp L1 and L2-3 at P21 (see Methods for details). We found that, compared to WT astrocytes, both the territory volumes and morphological complexity of LRRK2 G2019S astrocytes were significantly reduced (by ∼34%) (Figure 5A-C). However, this alteration of astrocyte morphology did not change the number of ALDH1L1+/SOX9+ cells (number of astrocytes) in the LRRK2 G2019S^ki/ki^ ACC and MOp (Figure S11B-D). Together, these results reveal that, *in vivo*, LRRK2 G2019S mutation reduces both astrocyte territory volume and morphological complexity without affecting astrocyte numbers.

**Figure 5:**
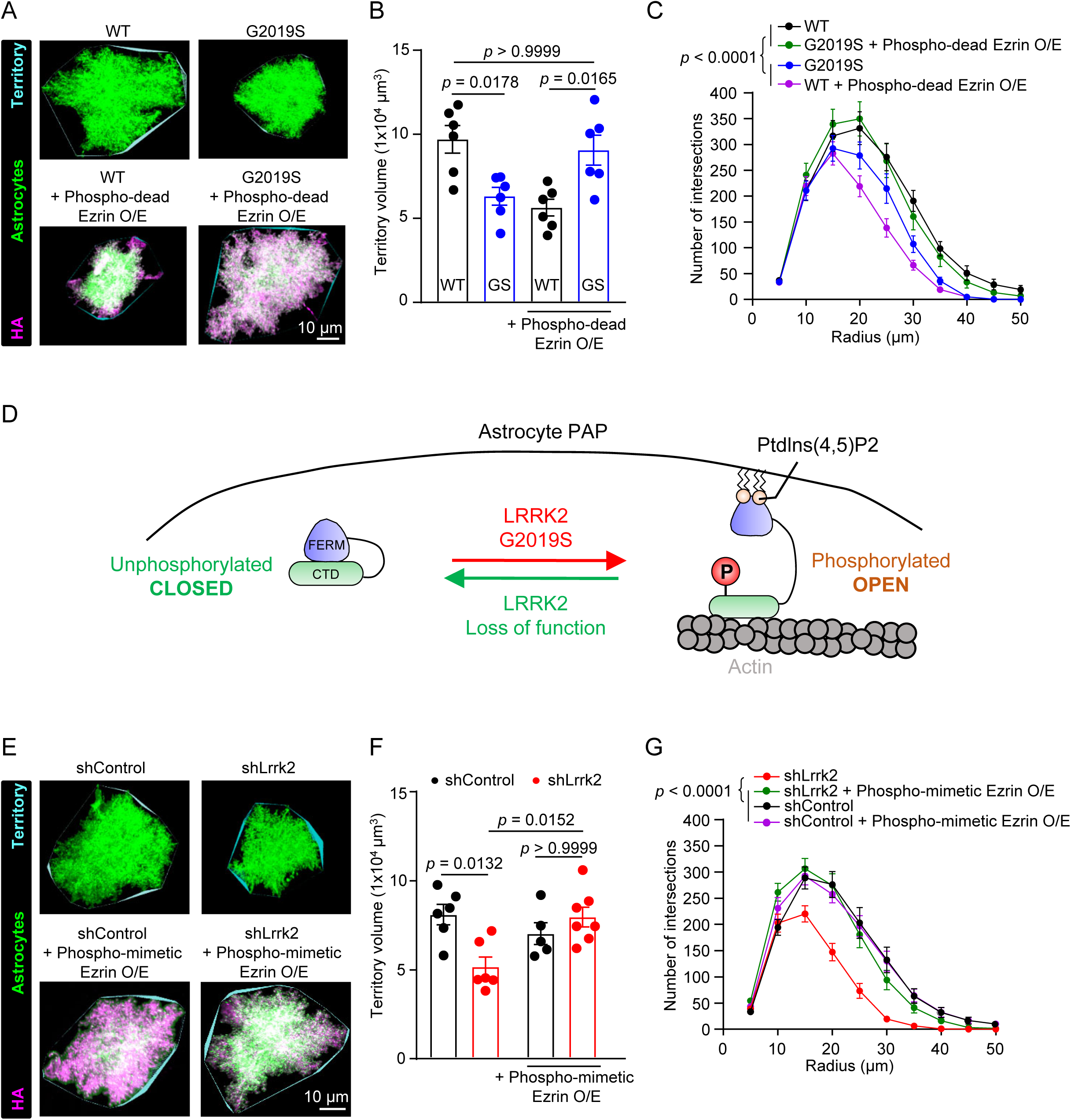
Astrocytic LRRK2 controls astrocyte morphology by balancing ERM phosphorylation levels *in vivo*. (A) Representative images of ACC and MOp L2-3 astrocytes in WT and LRRK2 G2019S^ki/ki^ (P21) expressing PB-mCherry-CAAX ± Phospho-dead EZRIN. Astrocyte territory in cyan. Scale bar, 10 μm. (B) Astrocyte territory volume (n = 16–18 astrocytes, 6 mice/group). One-way ANOVA [F(3, 20) = 7.987, p = 0.0011] with Bonferroni’s test showed significant differences: WT vs. LRRK2 G2019S^ki/ki^ (p = 0.0178), WT vs. WT + Phospho-dead EZRIN O/E (p = 0.0037), and WT + Phospho-dead EZRIN O/E vs. LRRK2 G2019S^ki/ki^ + Phospho-mimetic EZRIN O/E (p = 0.0165); other comparisons were not significant (p > 0.05). (C) Astrocyte branching complexity. N = 16–18 astrocytes (6 mice per group). Two-way ANOVA showed main effects of condition [F(2.505, 3305) = 158.4, *p* < 0.0001], radius [F(88, 1602) = 218.5, *p* < 0.0001], and their interaction [F(264, 3958) = 10.62, *p* < 0.0001]. Bonferroni’s test identified significant differences between groups (all *p* < 0.0001), except WT vs. LRRK2 G2019S^ki/ki^ + Phospho-dead EZRIN O/E (*p* = 0.5469). (D) Proposed model of Ezrin conformational transition and LRRK2 regulation in astrocyte morphogenesis. (E) Images of ACC and MOp L2-3 astrocytes expressing shControl-mCherry-CAAX or shLRRK2-mCherry-CAAX ± Phospho-mimetic EZRIN. Scale bar, 10 μm. (F) Astrocyte territory volume. n = 18–22 astrocytes (5–6 mice/group). One-way ANOVA [F(3, 19) = 5.47, *p* = 0.0070] with Bonferroni’s test showed significant differences between shControl and shLRRK2 (*p* = 0.0132), and shLRRK2 vs. shLRRK2 + Phospho-mimetic EZRIN O/E (*p* = 0.0152); no other comparisons were significant (*p* > 0.05). (G) Astrocyte branching complexity. n = 16–22 astrocytes. Two-way ANOVA revealed significant main effects of condition [F(2.435, 7008) = 225.0, *p* < 0.0001], radius [F(88, 8633) = 325.4, *p* < 0.0001], and their interaction [F(264, 8633) = 5.908, *p* < 0.0001]. Bonferroni’s test identified significant differences between most groups (all *p* < 0.0001) except shControl vs. shControl + Phospho-mimetic EZRIN O/E (*p* > 0.9999). Data are mean ± s.e.m.

Because phospho-dead Ezrin overexpression in astrocytes differentially impacts synapse densities and function in WT and LRRK2 G2019S^ki/ki^ mice (Figure 4), we wondered if overexpression of phospho-dead Ezrin would also impact astrocyte morphology differently between genotypes. We found that, in the ACC and MOp, the overexpression of phospho-dead Ezrin decreased the WT astrocyte territory volume and morphological complexity but restored LRRK2 G2019S^ki/ki^ astrocyte territory volume and complexity (Figure 5A-C). These results suggested that unphosphorylated Ezrin differentially regulates astrocyte morphology in WT versus LRRK2 G2019S^ki/ki^ astrocytes. We postulated that Ezrin’s conformational status may need to be tightly regulated for proper astrocyte morphogenesis (Figure 5D). In LRRK2 G2019S^ki/ki^ astrocytes, Ezrin is more likely to be in a phosphorylated and open state. This would deplete the presence of the closed unphosphorylated Ezrin. Such a distorted phospho-Ezrin balance may explain the morphological defects of LRRK2 G2019S^ki/ki^ astrocytes. To test this possibility, we sought to shift the balance in the opposing direction by reducing Ezrin phosphorylation through the knockdown of mouse Lrrk2. Similar to LRRK2 G2019S^ki/ki^ astrocytes, astrocytes transfected with shLrrk2 (∼90% *Lrrk2* reduction, Figure S12A-B) exhibited significantly smaller territory volumes (∼36% reduction) and reduced complexity (Figure 5E-G).

Previous studies suggested phosphorylation at T567 promotes Ezrin acquiring an open conformation by binding to phosphatidylinositol-4,5-bisphosphate (PIP2)[52]. In this conformation, the N-terminus interacts with membrane proteins, and the C-terminus is available to bind to the actin cytoskeleton[53, 54]. An Ezrin mutant in which the T567 residue is switched to aspartate (D) prefers Ezrin’s open conformation, thereby mimicking the phosphorylated state (Phospho-mimetic Ezrin, Figure S4C)[55, 56]. When we overexpressed phospho-mimetic Ezrin in astrocytes, we found that phospho-mimetic Ezrin restored the territory volume and morphological complexity of shLRRK2 astrocytes (Figure 5E-G) but had no effect on shControl astrocytes (Figure 5E-G). These results show that both the gain of LRRK2 kinase function (in LRRK2 G2019S^ki/ki^ mice) and the loss of LRRK2 function (shLRRK2) decrease astrocyte complexity and size and the manipulation of Ezrin conformational/phosphorylation states in opposing directions ( expression of phospho-dead or phospho-mimetic Ezrin respectively) can rescue these morphological phenotypes.

### LRRK2 G2019S mutation changes Ezrin interactome in astrocytes

To investigate how ERM hyperphosphorylation affects astrocyte morphology in LRRK2 G2019S^ki/ki^ astrocytes, we identified possible Ezrin interactors in WT and LRRK2 G2019S^ki/ki^ astrocytes by *in vivo* astrocyte-specific proximity labeling. We used two virally encoded astrocyte-specific biotin ligase probes: Ezrin fused with BioID (Astro-Ezrin-BioID) and soluble BioID with a nuclear export sequence (Astro-Cyto-BioID) (Figure 6A). These were delivered via intracortical AAV injections to P0-2 WT and LRRK2 G2019S^ki/ki^ mouse pups, followed by subcutaneous biotin injections starting at P18 for 3 days, and tissue collection at P21 (Figure 6B). Biotinylated proteins were detected in astrocytes using immunohistochemistry and immunoblotting for HA and Streptavidin (Figure S13).

**Figure 6:**
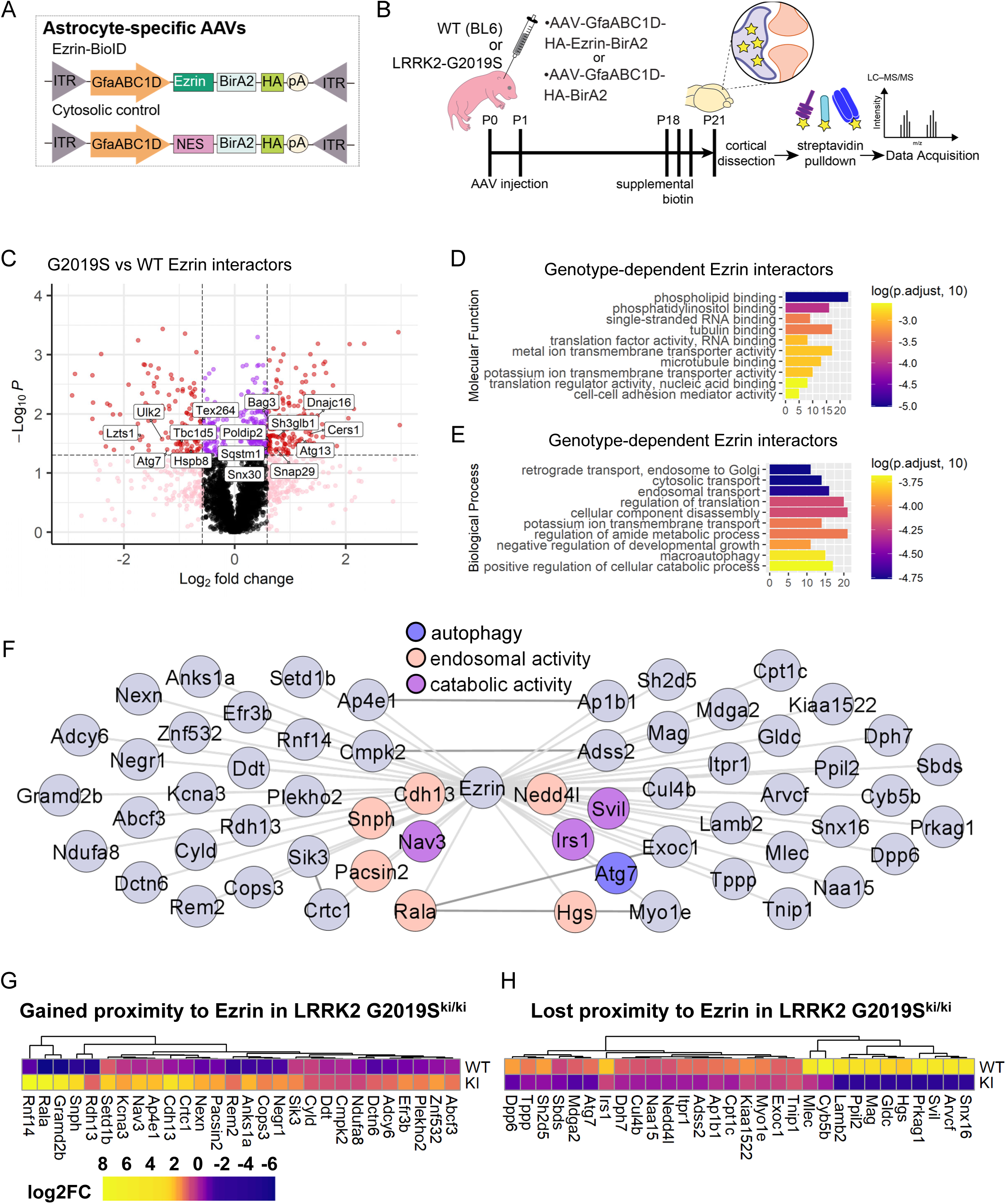
LRRK2 G2019S alters the interactome of astrocytic Ezrin *in vivo*. (A) astrocyte-specific AAVs (serotype PHP.eB) were used to express Ezrin or nonspecific cytosolic control. NES, nuclear export sequence; ITR, inverted terminal repeats; GfaABC1D truncated GFAP promoter; HA, hemagglutinin tag; pA, polyadenylation. (B) outline of the experimental paradigm. n = 3 biological replicates/construct/genotype (1 replicate = 2 animals per pooled sample). (C) Volcano plot showing the differential abundance of proteins detected by Astro-EZRIN-BioID in WT and LRRK2 G2019S^ki/ki^ cortices. (D-E) Bars show the top 10 most significant Gene Ontology (GO) terms, ordered by lowest adjusted p-value, for the proteins differentially detected by Astro-EZRIN-BioID in WT compared to LRRK2 G2019S^ki/ki^ (D) Molecular function (E) Biological Process. (F) The interaction network depicts 58 high-confidence proteins that gained or lost proximity to Ezrin in LRRK2 G2019S^ki/ki^ compared to WT mice. (G-H) heatmaps depict fold-change in abundance (Astro-EZRIN-BioID / Asto-Cyto-BioID) of proteins with high confidence changed proximity to Ezrin in LRRK2 G2019S^ki/ki^ astrocytes.

Affinity-purified proteins from cortices were analyzed using quantitative liquid chromatography-tandem mass spectrometry (LC-MS) (Figure 6B), identifying 30,815 unique peptides corresponding to 4,081 unique proteins (Supplemental Tables 2-4). Principle component analyses (PCA) of BioID results showed that distinct clustering between Cyto-BioID (control) and Ezrin-BioID (bait) samples, with further separation between WT or LRRK2 G2019S^ki/ki^ astrocytes (Figure S14A). Radixin and Moesin, known as Ezrin family proteins, were significantly enriched in the Ezrin-BioID sample compared to Cyto-BioID (Figure S14B)[57]. Comparative analysis revealed 344 proteins with significantly altered proximity to Ezrin in LRRK2 G2019S^ki/ki^ astrocytes (fold-change >1.5 or < -1.5, *p* < 0.05); 190 gained, and 154 lost Ezrin proximity (Figure 6C; Figure S14C-D; Supplemental Table 5). Gene Ontology (GO) analysis of these proteins showed enrichment in phospholipid binding, microtubule binding, and membrane transporter and cell-adhesion mediator activity were over-represented in differentially enriched proteins (Figure 6D; Supplemental Table 6). This aligns with our model that the LRRK2 G2019S mutation would shift Ezrin towards its phosphorylated/open state, promoting membrane interactions. Unexpectedly, GO-term analyses of biological processes highlighted enrichment in catabolic processes, macroautophagy, negative regulation of developmental growth, and intracellular transport (Figure 6E; Supplemental Table 7). These findings indicated that Ezrin plays a role in these biological processes, which are altered by the LRRK2 G2019S mutation.

Previous studies showed macroautophagy deficits in neurons with the LRRK2 G2019S mutation[58]. To assess global cytosolic changes, we analyzed the Astro-Cyto-BioID dataset across genotypes, identifying 195 differentially enriched proteins (78 gained, 117 lost in LRRK2 G2019S^ki/ki^; Figure S14E; Supplemental Table 8). GO-term analysis revealed an enrichment of proteins associated with phospholipid binding and vesicle trafficking (Figure S14F-G; Supplemental Tables 9-10). However, macroautophagy-related terms were not enriched, suggesting that the observed changes in Ezrin-BioID are specifically related to alterations in Ezrin function in the LRRK2 G2019S^ki/ki^ astrocytes.

BioID captures proteins in close proximity to the bait but does not guarantee direct interactions. To identify potential direct Ezrin interactors affected by LRRK2 G2019S, we compared protein abundance in Astro-Ezrin-BioID to Astro-Cyto-BioID within each genotype (Ezrin/Cyto fold -change > 1.5, *p* < 0.05, 346 proteins in WT and 335 proteins in LRRK2 G2019S^ki/ki^; Supplemental Tables 11-12). To generate high-confidence lists of interactors that gained or lost proximity to Ezrin in LRRK2 G2019S^ki/ki^ astrocytes, we set a second requirement for proteins with fold-change > 1.5 or < -1.5 between LRRK2 G2019S^ki/ki^ and WT Astro-Ezrin-BioID. Using these filters, we narrowed down our list to 28 as “gained proximity to Ezrin in LRRK2 G2019S astrocytes” and 30 as “lost proximity to Ezrin in LRRK2 G2019S astrocytes” (Figure 6F-H; Supplemental Table 13). Upregulated proteins, such as Rala, Cops3, and Anks1a, are involved in vesicular trafficking, cell signaling, and cytoskeletal regulation, aligning with enhanced ERM phosphorylation and their recruitment to the plasma membrane. In contrast, the downregulated proteins (e.g., Atg7, Cul4b, Myo1e, Irs1) suggest a disruption in intracellular trafficking and autophagy due to excessive ERM phosphorylation. Together, these data map Ezrin interactomes in WT and LRRK2 G2019S^ki/ki^ mouse cortical astrocytes *in vivo*, revealing changes in Ezrin interactions with proteins involved in catabolic processes and autophagy. This highlights a potential molecular mechanism linking LRRK2-Ezrin signaling to astrocyte morphogenesis.

### Atg7 is an Ezrin interactor that is required for proper astrocyte morphogenesis

Among high-confidence astrocytic Ezrin interactors, we identified autophagy-related gene 7 (Atg7), a crucial autophagy regulator, which was enriched (∼2.3-fold) in the Ezrin-BioID fraction of WT astrocytes but de-enriched (0.5-fold) in the Ezrin-BioID fraction of LRRK2 G2019S^ki/ki^ astrocytes compared to the Cyto-BioID. Atg7 is essential for autophagy initiation and has been implicated in neurological disorders[59],[60]. However, its role in astrocytes and its interaction with Ezrin is unclear. To explore this, we used AlphaFold-2.0 Multimer[61] to model the structures of Ezrin and Atg7 alone or in complex.

Atg7, a homodimer, has two globular domains: An N-terminal domain and a C-terminal domain that are separated by a short flexible linker. The C-terminal domain is further divided into the adenylation and extreme C-terminal domains (Figure 7A-B)[62]. Ezrin has an N-terminal FERM (four-point one, Ezrin, Radixin, Moesin) domain (subdivided into three subdomains-F1, F2, and F3), α-helical domain, and a C-terminal ERM Association domain (C-ERMAD) (Figure 7C)[56]. Upon phosphorylation of T567 and binding to PIP2, Ezrin adopts an open conformation (Figure 7D; see Methods for details of modeling)[63]. We then predicted the structure of the complex formed between the closed conformation of Ezrin and the homodimer of Atg7 (Figure 7E). Alpha-fold 2.0 multimer, in conjunction with CABS-Flex 2.0 (https://biocomp.chem.uw.edu.pl/CABSflex2, RRID: SCR_025674) and PyMol (http://www.pymol.org/, RRID: SCR_000305), were used to predict and assess conformational changes in Atg7 and Ezrin after binding each other (Figure 7F). When Ezrin binds Atg7, it is predicted to undergo drastic conformational changes in its α-helical domain, which is pulled towards the ERMAD and FERM domains, whereas the changes in the Atg7 dimer structure after Ezrin binding are minimal (Figure 7F).

**Figure 7:**
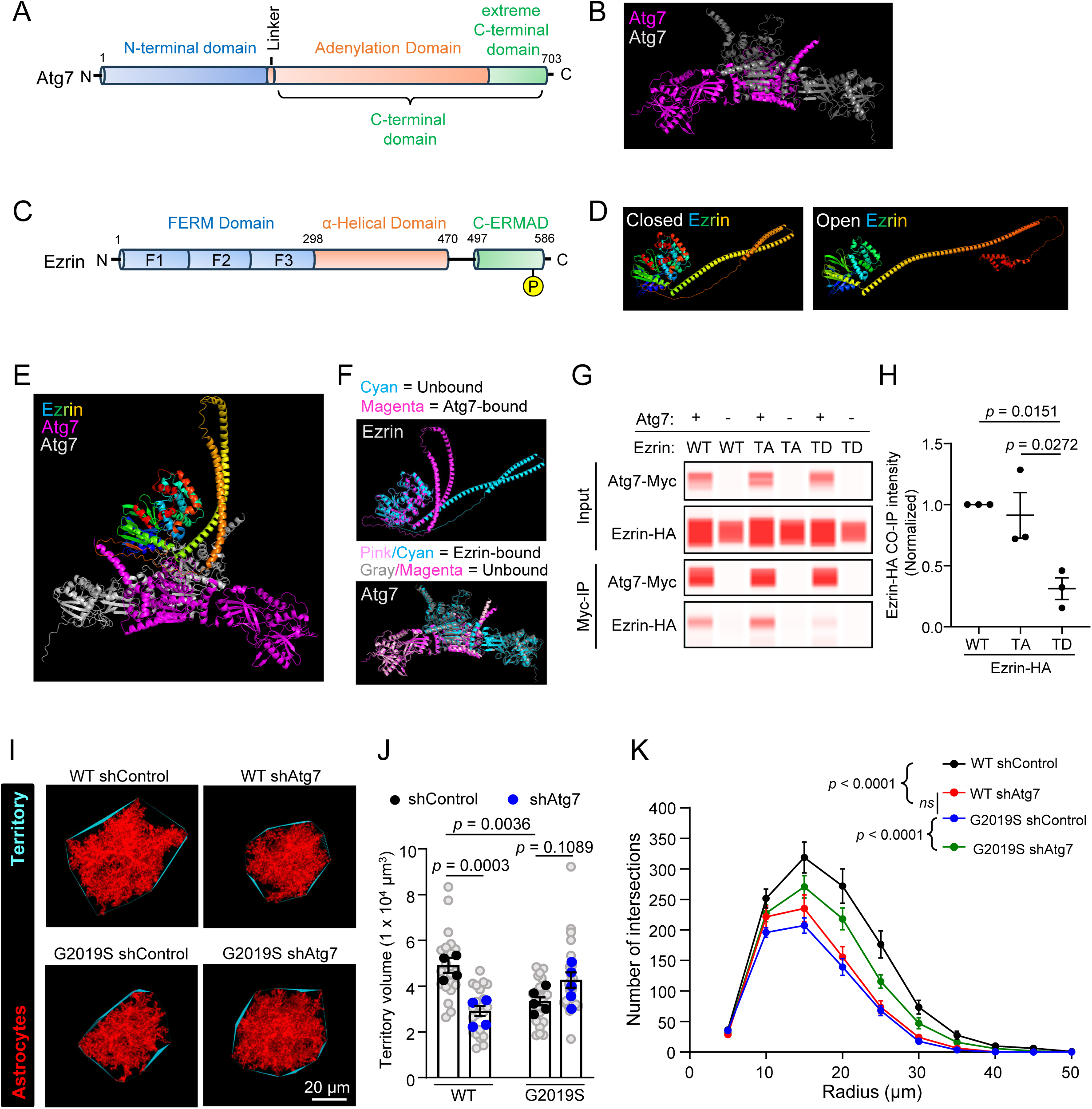
Interaction Between Atg7 and Ezrin Depends on Ezrin’s Phosphorylation State. (A) Schematic of Atg7 domains. (B) Predicted Atg7 homodimer model. (C) Schematic of Ezrin domains. (D) Predicted Ezrin structures in closed and open conformations. (E) Predicted Atg7-Ezrin interaction. (F) Structural changes in Ezrin and Atg7 upon binding. (G) Co-immunoprecipitation of Ezrin-HA by Atg7-Myc in HEK293T cells expressing WT, phospho-dead, or phospho-mimetic Ezrin. Ezrin was detected with anti-HA and Atg7 was detected with anti-Myc. (H) Quantification of (G). One-way ANOVA [F(2,6) = 9.932, *p* = 0.0125] with Tukey’s test: WT vs. phospho-mimetic Ezrin (*p* = 0.0151), phospho-dead vs. phospho-mimetic Ezrin (*p* = 0.0272), WT vs. phospho-dead Ezrin (*p* = 0.8658). n = 3 experiments. (I) Images of ACC and MOp L2-3 astrocytes at P21 expressing shControl or shAtg7-PB-mCherry-CAAX. Scale bar, 10 μm. (J) Quantification of astrocyte territories. Nested ANOVA [F(3,28) = 9.484, *p* = 0.0002] with Bonferroni tests: WT shControl vs. LRRK2 G2019S^ki/ki^ shControl (*p* = 0.0036), WT shControl vs. WT shAtg7 (*p* = 0.0003), WT shAtg7 vs. LRRK2 G2019S^ki/ki^ shAtg7 (*p* = 0.0109). n = 20–25 cells from 4–5 mice/group. (K) Astrocyte branching complexity. Two-way ANOVA revealed significant condition [F(2.324, 472.6) = 43.1, *p* < 0.05], radius [F(9, 250) = 301.8, *p* < 0.05], and interaction effects [F(27, 610) = 4.539, *p* < 0.05]. Bonferroni tests showed significant differences between groups, including WT shControl vs. LRRK2 G2019S^ki/ki^ shControl (*p* < 0.0001) and WT shControl vs. WT shAtg7 (*p* < 0.0001).

To validate the interaction between Ezrin and Atg7, we performed co-immunoprecipitation (Co-IP) in HEK293T cells overexpressing Myc-tagged Atg7 (Atg7-Myc) and HA-tagged WT Ezrin. To understand how the open or closed conformation of Ezrin would impact the Ezrin – Atg7 interaction, we also conducted Co-IPs with modulation of Ezrin phosphorylation levels: phospho-mimetic T567D (“TD”, open) and phospho-dead T567A (“TA”, closed) Ezrin. We found that WT Ezrin or phospho-dead Ezrin coimmunoprecipitated with Myc-Atg7, indicating that Atg7 interacts with WT Ezrin or phospho-dead Ezrin (Figure 7G). However, the interaction between Atg7 and phospho-mimetic Ezrin was significantly reduced (Figure 7H). These results support a phosphorylation-dependent interaction between Ezrin and Atg7, consistent with our BioID data showing reduced Atg7-Ezrin interaction in LRRK2 G2019S^ki/ki^ astrocytes.

To test if Atg7 is required for astrocyte morphology, we developed a short hairpin RNA (shRNA) that targets both rat and mouse *Atg7,* which effectively reduced *Atg7* mRNA in mouse cells (Figure S15A). Knockdown of Atg7 in rat astrocytes significantly reduced neuron-induced astrocyte morphogenesis *in vitro* in a neuron-astrocyte co-culture assay (Figure S15B-C). *In vivo*, Atg7 knockdown in WT astrocytes reduced astrocyte territory volume and morphological complexity, resembling the LRRK2 G2019S^ki/ki^ astrocytes transfected with shControl (Figure 7I-K). In LRRK2 G2019S^ki/ki^ astrocytes, Atg7 knockdown improved morphological complexity but did not further reduce territory volume (Figure 7J-K). These findings highlight Atg7’s role in regulating astrocyte morphology and suggest its dysfunction in LRRK2 G2019S^ki/ki^ astrocytes contributes to altered morphology.

## DISCUSSION

Here, we describe a previously unknown role for LRRK2, a PD-linked gene, in controlling astrocyte morphological complexity by regulating the phosphorylation state of ERM proteins. In the presence of LRRK2 G2019S mutation, we found enhanced ERM phosphorylation in human and mouse cortical astrocytes. ERM hyperphosphorylation in the LRRK2 G2019S background is specific to a subset of the ACC and MOp astrocytes of juvenile and adult mice. Previous studies suggested a direct interaction between LRRK2 and ERM proteins[18] and that ERM proteins are substrates of LRRK2[23, 24]. However, whether the enhanced ERM phosphorylation in LRRK2 G2019S astrocytes is due to a direct effect of LRRK2 kinase is unclear. Several other kinases, such as lymphocyte-oriented kinase, STE20-like protein kinase (SLK), and Rho kinase, were found to be involved in ERM phosphorylation[64], suggesting the enhanced ERM phosphorylation we observed in LRRK2 G2019S brains may be an indirect effect of the LRRK2 G2019S mutation on the activities of these kinases.

### How does ERM phosphorylation control astrocyte morphology and synaptogenic functions?

The canonical function of ERM proteins is linking the actin cytoskeleton with the plasma membrane, thus allowing cellular process outgrowth and morphogenesis[53]. Our findings suggest that disruptions in the interaction between Ezrin and a core autophagy-related protein, Atg7, contribute to the altered astrocyte morphology observed in LRRK2 G2019S^ki/ki^ mice. *In vivo* Ezrin interactome in astrocytes included several other proteins related to autophagy, which changed their proximity to Ezrin in the LRRK2 G2019S^ki/ki^ astrocytes, compared to WT astrocytes. These observations indicate that LRRK2 G2019S-induced ERM hyperphosphorylation may impair astrocyte autophagy and cause astrocyte dysfunction.

Another intriguing observation is that, *in vivo*, knocking down Atg7 in WT astrocytes reduces astrocyte morphological complexity. Atg7 is a homodimeric E1-like enzyme essential for autophagy, which maintains cellular homeostasis by degrading and recycling damaged organelles and proteins[59]. Autophagic dysregulation is linked to neurodegenerative diseases, including those associated with PD-related LRRK2 mutations[40, 65],[58]. However, the role of autophagy in astrocyte morphology and synaptogenic functions is not well understood, nor is the impact of LRRK2 G2019S on autophagy in astrocytes. It is possible that Atg7 depletion disrupts autophagic flux, causing the accumulation of damaged components that compromise astrocyte structure and function.

Our data also show that Atg7 knockdown in LRRK2 G2019S astrocytes does not worsen their morphological defects. The G2019S mutation enhances LRRK2’s kinase activity, leading to aberrant phosphorylation of Rab proteins and trapping them on the membrane surface[66]. Our studies revealed that Atg7/Ezrin interaction requires the unphosphorylated form of Ezrin. It is plausible that this interaction, which is impaired in LRRK2 G2019S astrocytes, diminished astrocytic Atg7 function. Therefore, additional knockdown of Atg7 in LRRK2 G2019S astrocytes does not further reduce their size and complexity. Instead, LRRK2 G2019S astrocytes may try to compensate by upregulating catabolic pathways like ubiquitination and proteasomal degradation. These mechanisms are crucial for clearing misfolded or damaged proteins in the absence of functional endolysosomal function and may help mitigate the loss of autophagic flux[67, 68]. Our Ezrin BioID dataset also showed that Nedd4l, an E3 ubiquitin ligase[69], lost proximity to Ezrin in LRRK2 G2019S astrocytes (Figure 6).

Astrocytes regulate synapse formation and function through cell adhesion molecules and secreted proteins that signal to neurons[6]. Synaptogenic proteins like Hevin and thrombospondins shape cortical excitatory circuits[70],[71]. ERM proteins are known to link the actin cytoskeleton with the plasma membrane, supporting cellular process outgrowth and morphogenesis[63, 72, 73], but their role in secretion is less understood. Phosphorylation of ERM proteins may refine secretory mechanisms in astrocytes or regulate synapse elimination via phagocytosis. In agreement with this possibility, a recent study found that synapses are more likely to be eliminated if phospho-ERM proteins are localized to peri-synaptic astrocyte processes in ALS patient brains and mouse models[74]. Future research exploring the relationships between LRRK2, ERM, and autophagy in astrocytes will help clarify these processes and their impact on brain region-specific excitatory and inhibitory synapse phenotypes.

### Implications for Therapeutic Development

LRRK2 G2019S^ki/ki^ mice do not recapitulate motor symptoms of PD because there is no detectable dopaminergic neuron loss in these mice[41, 75]. However, these mice may serve as good models for the non-motor (prodromal) PD symptoms seen in LRRK2 G2019S carriers[76]. These non-motor symptoms, which arise long before motor dysfunction in PD, indicate aberrant synaptic connectivity within multiple brain circuits, including the cortex, long before the death of dopaminergic neurons[77]. LRRK2 G2019S^ki/ki^ mice display early circuit dysfunction and non-motor symptoms akin to prodromal PD. Previous studies showed that LRRK2 G2019S^ki/ki^ mice have deficits in attention and goal-directed learning and differential response to stress[34, 78, 79]. Our findings showing synaptic deficits in the ACC of LRRK2 G2019S^ki/ki^ mice (Figure 2-3) fit well with these studies because the ACC is a brain region that controls attention, goal-directed actions, and mood[80, 81]. Furthermore, we observed synaptic deficits in the ACC of LRRK2 G2019S^ki/ki^ mice concurrently with astrocyte morphological deficits located in the ACC and MOp.

Why do ERM phosphorylation and synaptic deficits happen in specific brain regions of LRRK2 G2019S^ki/ki^ mice? Cortical astrocytes are heterogeneous in their gene expression, morphology, and function[82, 83]. Interestingly, cortical astrocyte heterogeneity is controlled by layer-specific neuron heterogeneity[82, 83]. It is plausible that neurons and astrocytes in different cortical regions have varying levels of LRRK2 and diverse susceptibilities to LRRK2 mutations. The ACC and MOp may highly depend on the LRRK2-ERM pathway to maintain synaptic integrity and astrocyte function. A recent single-nucleus RNA sequencing in the striatum revealed that the differential expression level of LRRK2 is correlated with changes in the ciliogenesis of different cholinergic neurons[84], suggesting that LRRK2 transcriptional states influence cell function in a cell-type-specific manner. Future studies investigating the impacts of LRRK2 G2019S on cortical neurons and astrocytes’ transcriptional profiles may help explain why specific brain regions and astrocyte populations are susceptible to the LRRK2 G2019S mutation.

In sum, our findings reveal that LRRK2 G2019S mutation-mediated ERM hyperphosphorylation in astrocytes causes morphological dysfunction in cortical astrocytes. Our results suggest that this astrocytic dysfunction could directly contribute to excitatory synaptic deficits observed in the ACC of the LRRK2 G2019S mice. Restoring astrocyte function by inhibiting ERM hyperphosphorylation, even post-development, was sufficient to rectify excitatory synaptic pathology. Future studies aiming to identify the molecular mechanisms by which LRRK2 controls ERM phosphorylation and the specific phosphatases involved could lead to new therapeutic targets for PD.

### Limitations of the study

In our study, we found that a subset of astrocytes in the ACC and MOp were responsive to LRRK2 kinase activity. We also pinpoint that the balance of ERM phosphorylation plays a crucial role in regulating astrocyte morphology in the ACC and MOp. However, astrocytes display molecular, morphological, and functional heterogeneities[85]. We do not yet know why these astrocytes in the ACC and MOp would be sensitive to LRRK2 kinase activity and how these astrocytes are unique compared to other astrocytes in terms of their transcriptional and proteomic profiles. Another future direction would be to determine how the ERM phosphorylation balance modulates astrocyte morphological changes and synapse changes. ERM is a crucial player in linking F-actin to the cell membrane and is important for regulating membrane tension. Actin cytoskeleton dynamics are important for astrocyte morphogenesis[86]. Future single-cell multi-omics studies and experiments understanding how ERM phosphorylation alters actin cytoskeleton dynamics in astrocytes will help us address these current limitations. In addition, the difference in GFAP levels between human PD patients and LRRK2 G2019S^ki/ki^ mice reflects the complexity of human PD pathology, the limitations of single-gene mouse models, and potential species-specific differences in astrocyte reactivity. This discrepancy highlights the challenges in modeling human neurodegenerative diseases in animal models and underscores the need for caution when translating findings from animal studies to human disease. Furthermore, in our study, we used PALE to label astrocytes. While PALE is a powerful tool that allows us to label a sparse population of astrocytes in the brain and study their morphological complexity in detail, it has limitations in labeling astrocytes in deep brain regions like the striatum and substantia pars compacta. Future studies using systemic approaches may overcome this technical limitation and clarify whether and how LRRK2 G2019S affects astrocyte morphology and function in additional brain regions.

## MATERIALS AND METHODS

### KEY RESOURCES TABLE

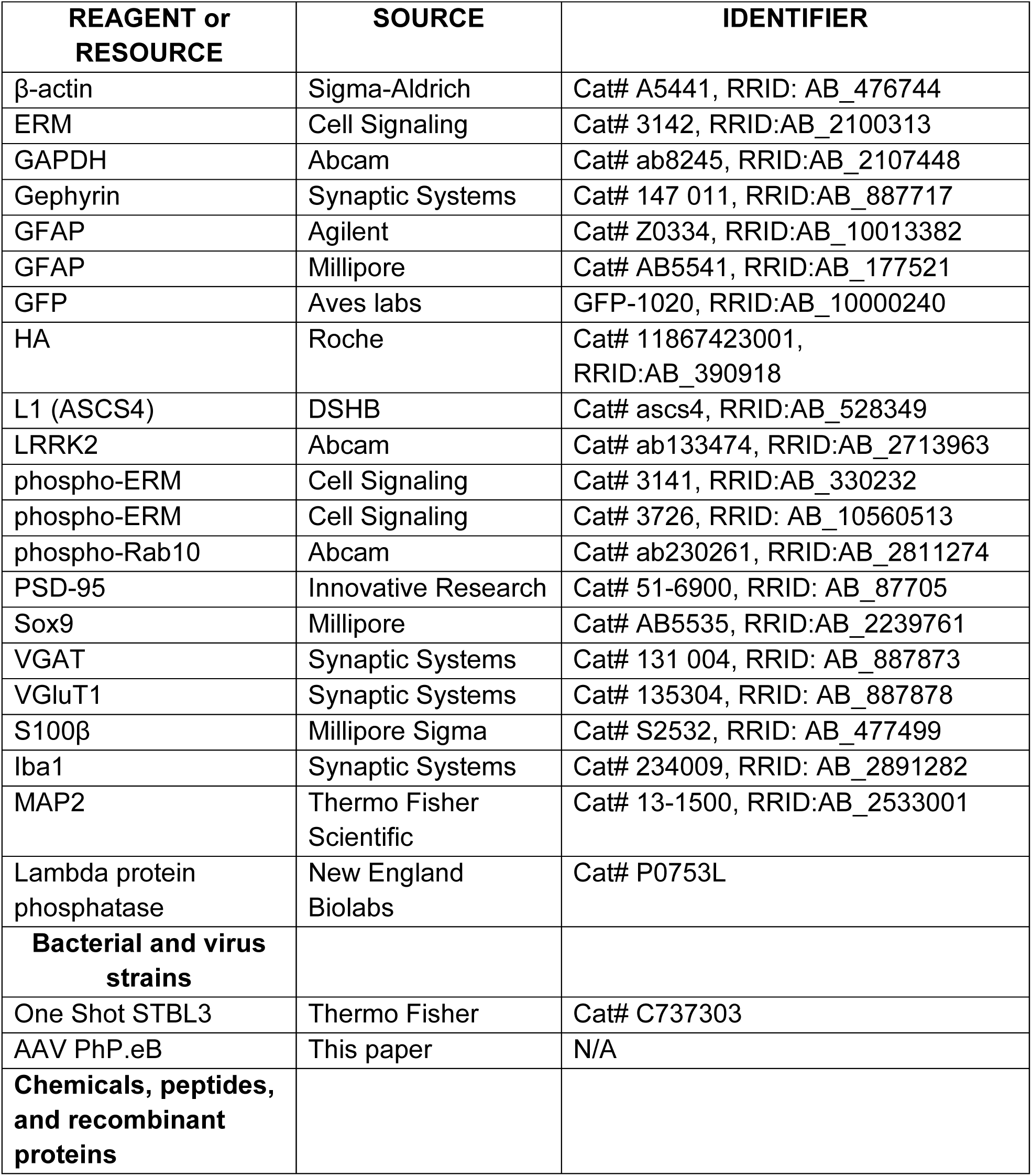
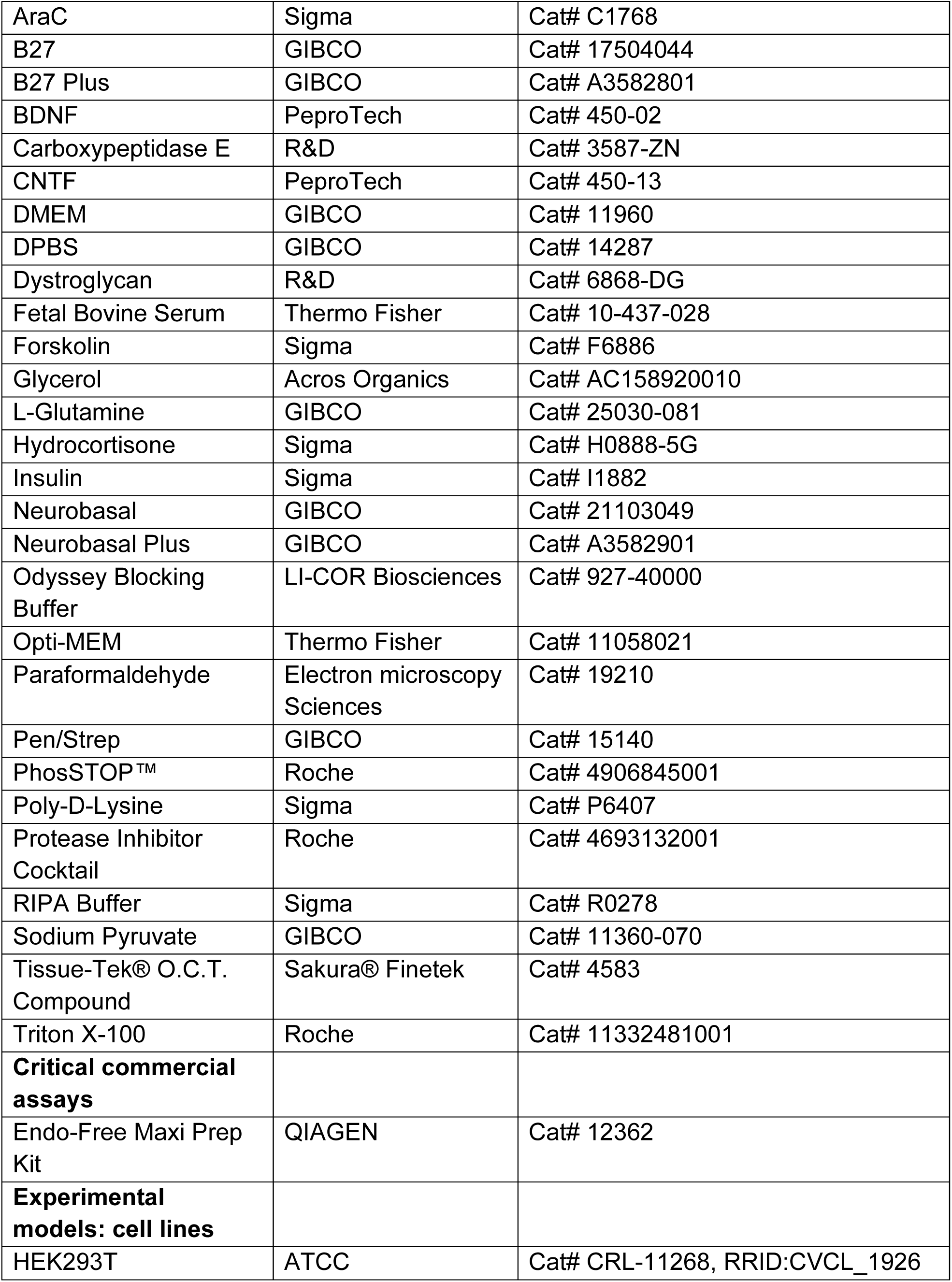
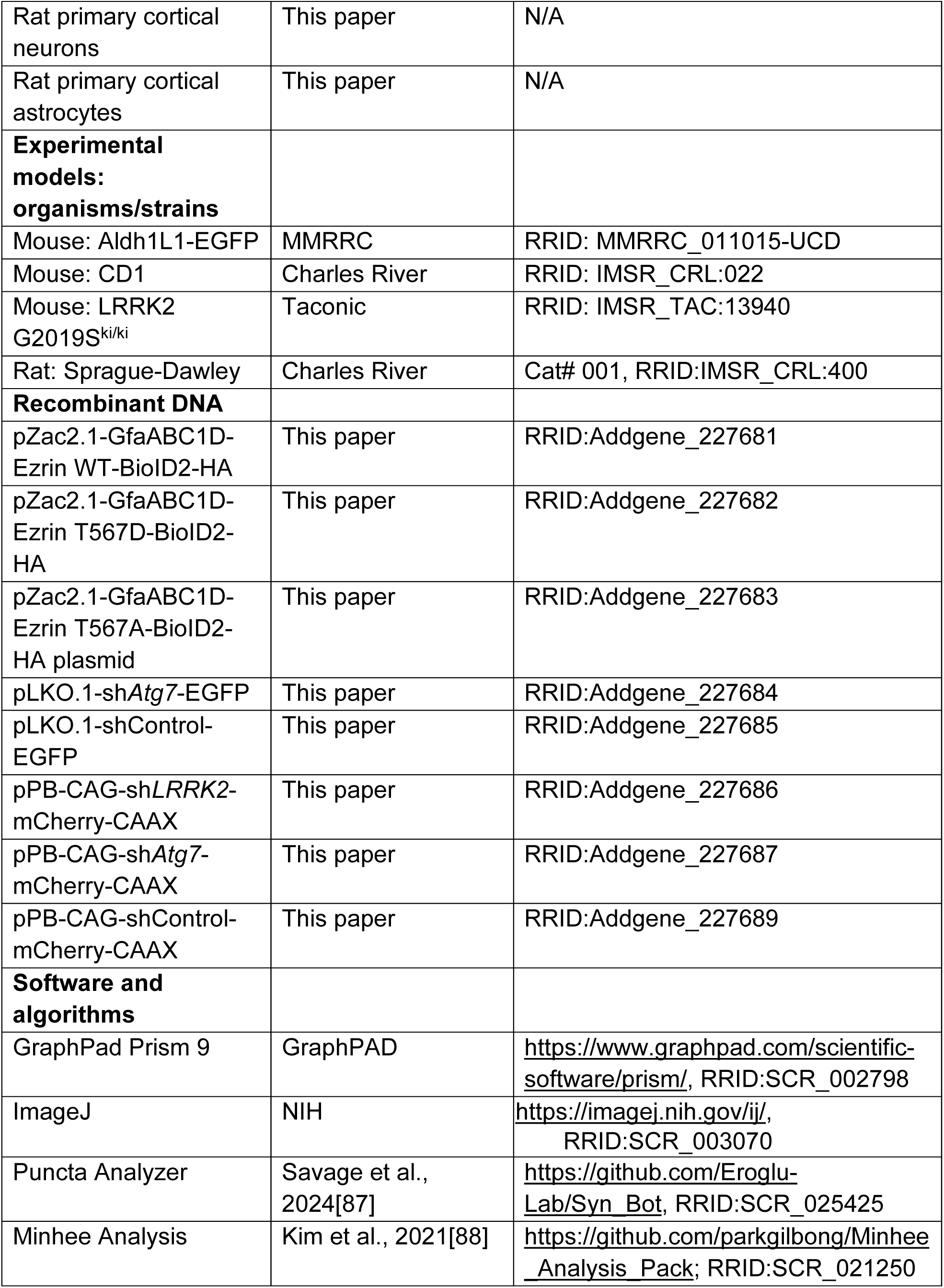
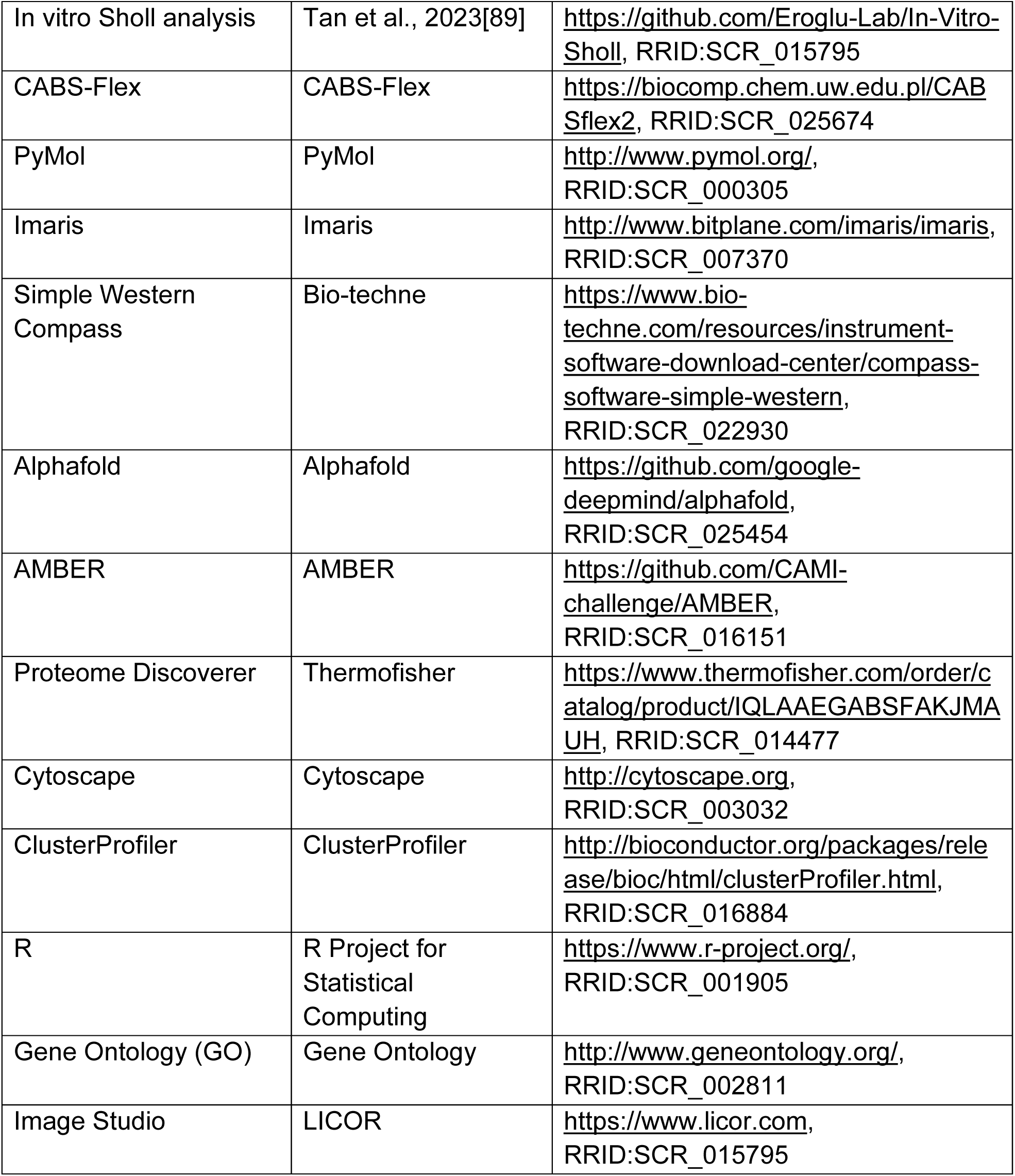

### RESOURCE AVAILABILITY

#### Lead contact

Further information and requests for resources and reagents should be directed to and will be fulfilled by the lead contacts, Shiyi Wang and Cagla Eroglu.

#### Materials availability

The reagents generated in this study are available without restriction.

#### Data availability

The data generated during this study are available at the link https://zenodo.org/records/15282825.

The proteomic data is uploaded to the repository ProteomeXchange via the PRIDE database (https://www.ebi.ac.uk/pride/), project accession number: PXD062312.

#### Code availability

The code used to analyze Sholl data is available via the link https://github.com/Eroglu-Lab/Sholl_Macro.

#### Protocol availability

The protocols used for this study are available at the link https://www.protocols.io/researchers/shiyi-wang1/publications

### EXPERIMENTAL MODELS AND SUBJECT DETAILS

#### Animals

All mice and rats were used in accordance with the Institutional Animal Care and Use Committee (IACUC) and the Duke Division of Laboratory Animal Resources (DLAR) oversight (IACUC Protocol Numbers A147-17-06 and A117-20-05). All mice and rats were housed under typical day/night conditions of 12-hour cycles. LRRK2 G2019S^ki/ki^ mice (RRID: IMSR_TAC:13940) were obtained through Taconic. This model was generated on the C57BL6/NTac background. It has been maintained on that pure background in Taconic. Aldh1l1-EGFP (RRID: MMRRC_011015-UCD) mice were obtained through MMRRC and backcrossed on a C57BL/6J background. Wild-type CD1 mice used for PALE were purchased through Charles River Laboratories (RRID: IMSR_CRL:022).

Mice and rats were used for experiments as specified in the text and figure legends. For all experiments, age, and sex-matched mice were randomly assigned to experimental groups based on genotypes. Mice of both sexes were included in the analysis, and we did not observe any influence or association of sex on the experimental outcomes.

#### Primary cortical neuron isolation and culture

Purified (glia-free) rat cortical neurons were prepared as described previously[36]. Briefly, cortices from P1 rat pups of both sexes (Sprague Dawley, Charles River Laboratories, SD-001) were micro-dissected, digested in papain (∼7.5 units/ml) at 33°C for 45 minutes, triturated in low and high ovomucoid solutions, resuspended in panning buffer (DPBS (GIBCO 14287) supplemented with BSA and insulin) and passed through a 20 μm mesh filter (Elko Filtering 03-20/14). Filtered cells were incubated on negative panning dishes coated with Bandeiraea Simplicifolia Lectin 1 (x2), followed by goat anti-mouse IgG+IgM (H+L) (Jackson ImmunoResearch 115-005-044), and goat anti-rat IgG+IgM (H+L) (Jackson ImmunoResearch 112-005-044) antibodies, then incubated on positive panning dishes coated with mouse anti-L1 (ASCS4, Developmental Studies Hybridoma Bank, Univ. Iowa) to bind cortical neurons. Adherent cells were collected by forceful pipetting with a P1000 pipette. Isolated neurons were pelleted (11 minutes at 200 g) and resuspended in serum-free neuron growth media (NGM; Neurobasal, B27 supplement, 2 mM L-Glutamine, 100 U/ml Pen/Strep, 1 mM sodium pyruvate, 4.2 μg/ml Forskolin, 50 ng/mL BDNF, and 10 ng/mL CNTF). 70,000 neurons were plated onto 12 mm glass coverslips coated with 10 μg/ml poly-D-lysine (PDL, Sigma P6407) and 2 μg/ml laminin and incubated at 37°C in 10% CO_2_. On day *in-vitro* (DIV) 2, half of the media was replaced with NGM Plus (Neurobasal Plus, B27 Plus, 100 U/mL Pen/Strep, 1 mM sodium pyruvate, 4.2 μg/ml Forskolin, 50 ng/ml, BDNF, and 10 ng/ml CNTF) and AraC (10 μM) was added to stop the growth of proliferating contaminating cells. On DIV 3, all the media was replaced with NGM Plus. In experiments involving lentivirus infection, 100 μl of supernatant containing lentivirus plus polybrene (1 μg/ml) was added to the AraC NGM mixture on DIV 2 and completely washed out on DIV 3 and replaced with NGM Plus containing 100 ng/ml BDNF. Neurons were fed on DIV 6 and DIV 9 by replacing half of the media with NGM Plus.

#### Primary cortical astrocyte isolation and culture

##### Rat astrocyte isolation and culture

Rat cortical astrocytes were prepared as described previously[36]. P1 rat cortices from both sexes were micro-dissected, papain digested, triturated in low and high ovomucoid solutions, and resuspended in astrocyte growth media (AGM: DMEM (GIBCO 11960), 10% FBS, 10 μM, hydrocortisone, 100 U/ml Pen/Strep, 2 mM L-Glutamine, 5 μg/ml Insulin, 1 mM Na Pyruvate, 5 μg/ml N-Acetyl-L-cysteine). Between 15-20 million cells were plated on 75 mm^2^ flasks (non-ventilated cap) coated with poly-D-lysine and incubated at 37°C in 10% CO_2_. On DIV 3, the removal of non-astrocyte cells was performed by forcefully shaking closed flasks by hand for 10-15 s until only an adherent monolayer of astrocytes remained. AraC was added to the media from DIV 5 to DIV 7 to eliminate contaminating fibroblasts. On DIV 7, astrocytes were trypsinized (0.05% Trypsin-EDTA) and plated into 12-well or 6-well dishes. On DIV 8, cultured rat astrocytes were transfected with shRNA and/or expression plasmids using Lipofectamine LTX with Plus Reagent (Thermo Scientific) per the manufacturer’s protocol. Briefly, 1 μg (12-well) or 2 μg (6-well) total DNA was diluted in Opti-MEM containing Plus Reagent, mixed with Opti-MEM containing LTX (1:2 DNA to LTX), and incubated for 30 minutes. The transfection solution was added to astrocyte cultures and incubated at 37°C for 3 hours. On DIV 10, astrocytes were trypsinized, resuspended in NGM plus, plated (20,000 cells per well) onto DIV 10 neurons, and co-cultured for 48 hours.

##### shRNA plasmids

pLKO.1 Puro plasmids containing shRNA (pLKO.1-shRNA) against mouse/rat *Lrrk2* (sh*LRRK2*: TRCN0000322193; GGCCGAGTTGTGGATCATATT) and mouse/rat *Atg7* (sh*Atg7*: TRCN0000092163; TATTATTGAGTTCAGAGCTGG), were obtained from the RNAi Consortium (TRC) via Dharmacon. Two scrambled shRNA sequences were generated (GTTGCTGAATGGCGGATCTAT; GTTGCGGTTATGAATAGTACT) and cloned into the pLKO.1 TRC cloning vector[90] according to Addgene protocols (https://www.addgene.org/protocols/plko/). To generate pLKO.1 shRNA plasmids that express EGFP (pLKO.1-shRNA-EGFP), CAG-EGFP was removed from pLenLox-shNL1-CAG-EGFP[91] and inserted between Kpn1 and SpeI sites in pLKO.1 Puro, replacing the puromycin resistance gene. pLKO.1 shRNA mCherry plasmids were generated by replacing EGFP with mCherry between KpnI and NheI sites.

##### PiggyBac plasmids

pPB-CAG-EGFP (RRID:Addgene_40973) and pGLAST-PBase were a gift from Dr. Joseph Loturco[92]. To generate pPB-CAG-mCherry-CAAX, mCherry-CAAX was inserted between XmaI and NotI restrictions sites to replace EGFP. To insert the hU6 promoter and shRNA in pPB-CAG-mCherry-CAAX, a DNA fragment containing hU6 and shRNA was amplified from pLKO.1-shRNA using Phusion High-Fidelity DNA Polymerase (NEB) with primers that introduced SpeI restriction sites (Forward Primer: GGACTAGTCAGGCCCGAAGGAATAGAAG; Reverse Primer: GGACTAGTGCCAAAGTGGATCTCTGCTG). PCR products were purified, digested with SpeI, and ligated into pPB-CAG-mCherry-CAAX at the SpeI restriction site. An analytical digest with EcoRI followed by sequencing was used to confirm the orientation of the inserted DNA fragment.

##### Ezrin plasmids

pZac2.1-GfaABC1D-Lck-GCaMP6f was a gift from Dr. Baljit Khakh (Addgene plasmid #52924, RRID:Addgene_52924). pZac2.1-GfaABC1D-BioID2-HA was generated by PCR of BioID2 from pAAV-hSyn-BioID2-Linker-Synapsin1a-HA[93] (primers Fwd: 5’- ctagcctcgagaattcaccatgttcaaaaatcttatttg-3’, Rev: 5’-ccgggtcgactctagatgcgtaatccggtacatcg-3’) and insertion into the EcoRI and XbaI restriction sites of pZac2.1-GfaABC1D-Lck-GCaMP6f using In-Fusion cloning (TaKaRa). pHJ421(pEGFP-Ezrin WT) and pHJ423 (pEGFP-Ezrin T567D) were a gift from Stephen Shaw (Addgene plasmid # 20680 and # 20681; RRID:Addgene_20680, RRID:Addgene_20681). pZac2.1-GfaABC1D-Ezrin WT-BioID2-HA (RRID:Addgene_227681) and pZac2.1-GfaABC1D-Ezrin T567D-BioID2-HA (RRID:Addgene_227682) were generated by PCR of Ezrin from pHJ421 or pHJ423 (primers Fw: 5’-ctagcctcgagaattcaccatgccgaaaccaatca-3’, Rv: 5’- tgaacatggtgaattccgacagggcctcgaactcg-3’) and insertion into the EcoRI restriction sites of pZac2.1-GfaABC1D-BioID2, respectively. To generate pZac2.1-GfaABC1D-Ezrin T567A-BioID2-HA plasmid, Q5® Site-Directed Mutagenesis Kit (NEB) was used with mutagenesis primers (Fwd: CAAGTACAAGGCGCTGCGGCAGA; Rev: TCCCGGCCTTGCCTCATG)

##### Lentivirus production and transduction

Lentiviruses containing shRNA targeting vectors were produced to test the knockdown efficiency of shRNA constructs in cultured primary astrocytes or to bulk transduce neurons with shRNA and GFP. To produce lentivirus, HEK293T cells were transfected with a pLKO.1 shRNA Puro targeting plasmid (for astrocyte transduction), an envelope plasmid (VSVG) (RRID:Addgene_12259), and a packaging plasmid (dR8.91) using X-tremeGENE (Roche). One day after transfection, the media was replaced with AGM (for astrocyte transduction), and media containing lentivirus was collected on days 2 and 3 post-transfection. To assess the knockdown efficiency of shRNAs in astrocytes, rat primary astrocytes at DIV 7 were plated in 6-well dishes in 2 ml of AGM. On DIV 8, 1 ml of AGM was removed, and 500 μl of fresh AGM was added along with 500 μl of lentivirus-containing media and 1 μg/ml polybrene. Cultured astrocytes were treated with puromycin (1 μg/ml) from DIV 10-15 to select for transduced cells. Cultured astrocytes were lysed at DIV 15 for protein extraction and Western blot analysis.

##### Immunocytochemistry

Astrocyte-neuron co-cultures on glass coverslips were fixed on DIV 12 with warm 4% PFA for 7 minutes, washed 3 times with PBS, blocked in a blocking buffer containing 50% normal goat serum (NGS) and 0.4% Triton X-100 for 30 minutes, and washed in PBS. Samples were then incubated overnight at 4°C in primary antibodies diluted in blocking buffer containing 10% NGS, washed with PBS, incubated in Alexa Fluor conjugated secondary antibodies (Life Technologies) for 2 hours at room temperature, and washed again in PBS. Coverslips were mounted onto glass slides (VWR Scientific) with Vectashield mounting media containing DAPI (Vector Labs), sealed with nail polish, and imaged on an AxioImager M1 (Zeiss) fluorescence microscope. Images of healthy astrocytes with strong expression of fluorescent markers that did not overlap with other fluorescent astrocytes were acquired at 40x magnification in red, green, and/or DAPI channels using a CCD camera. Astrocyte morphological complexity was analyzed in FIJI using the Sholl analysis plugin[94], as described previously[36] (https://github.com/Eroglu-Lab/In-Vitro-Sholl, RRID: SCR_022758). Statistical analyses were conducted using a custom code in R (https://github.com/Eroglu-Lab/In-Vitro-Sholl, RRID: SCR_022758). A mixed-effect model with multiple comparisons made using the Tukey post-test was used for Sholl analysis to account for the variability per experiment as a random effect to evaluate differences between the conditions. The exact number of independent experiments and the exact number of cells analyzed are indicated in the figure legend for each experiment. To ensure the health of astrocyte-neuron co-cultures, the peak for the number of astrocyte intersections must be greater than or equal to 25 in the control condition. We imaged non-overlapping astrocytes that contained a single nucleus (DAPI stain) and expressed consistent fluorescent markers and Ezrin constructs according to the experimental conditions.

##### Postnatal astrocyte labeling by electroporation (PALE)

Late P0/early P1 mice were sedated by hypothermia until anesthetized, and 1 μl of plasmid DNA mixed with Fast Green Dye was injected into the lateral ventricle of one hemisphere using a pulled glass pipette (Drummond). For shRNA knockdown experiments in wild-type CD1 mice, the 1 μl of DNA contained 1 μg of pGLAST-PBase and 1 μg of pPB-shRNA-mCherryCAAX was injected. To label astrocytes in WT and LRRK2 G2019S^ki/ki^ mice, the 1 μl of DNA contained 1 μg of pGLAST-PBase and 1 μg of pPB-mCherry-CAAX was injected per mouse.

For PALE-mediated overexpression of phospho-mimetic Ezrin in shRNA knockdown experiments, 0.5 μg pGLAST-PBase, 0.5 μg pPB-shRNA-mCherryCAAX, and 1 μg pZac2.1-GfaABC1D-Ezrin T567D-BioID2-HA were injected in a total volume of 1 μl. For phospho-dead Ezrin overexpression in WT and LRRK2 G2019S^ki/ki^ mice, 0.5 μg pGLAST-PBase, 0.5 μg of pZac2.1-gfaABC1D-mCherry-CAAX and 1 μg of pZac2.1-GfaABC1D-Ezrin T567A-BioID2-HA were injected in a total volume of 1 μl. Following DNA injection, electrodes were oriented with the positive terminal above the frontal cortex and the negative terminal below the chin, and 5 discrete 50 ms pulses of 100 V spaced 950 ms apart were applied. Pups were recovered on a heating pad, returned to their home cage, and monitored until collection at P21. The number of mice used for each experiment is indicated in the figure legends. All animals appeared healthy at the time of collection. Brain sections were examined for the presence of electroporated cells before staining.

##### Adeno-associated virus (AAV) production and administration

Purified AAVs were produced as described previously[95]. Briefly, HEK293T cells were transfected with pAd-DELTA F6, serotype plasmid AAV PHP.eB (RRID: Addgene_103005), and AAV plasmid (pZac2.1-GfaABC1D-Ezrin-BioID2-HA; RRID:Addgene_227681 or pZac2.1-GfaABC1D-Ezrin T567A-BioID2-HA; RRID:Addgene_227683). Three days after transfection, cells were collected in 15 mM NaCl, 5 mM Tris-HCl, pH 8.5, and lysed with repeat freeze-thaw cycles followed by treatment with Benzonase (Novagen 70664) at 37°C for 30 minutes. Lysed cells were pelleted by centrifugation, and the supernatant, containing AAVs, was applied to an Optiprep density gradient (Sigma D1556, 15%, 25%, 40%, and 60%) and centrifuged at 67,000 rpm using a Beckman Ti-70 rotor for 1 hour. The AAV-enriched fraction was isolated from between 40% and 60% iodixanol solution and concentrated by repeated washes with sterile PBS in an Amicon Ultra-15 filtration unit (NMWL: 100 kDa, Millipore UFC910008) to a final volume of ∼100 μl and aliquoted for storage at −80°C. 9-week-old WT or LRRK2 G2019S^ki/ki^ mice placed in a stereotaxic frame were anesthetized through inhalation of 1.5% isofluorane gas. 10 μl of purified AAVs having a titer of ∼1 x 10^12^ GC/ml was introduced into the mouse brain intravenously by injection into the retro-orbital sinus. After 3 weeks at 12-week-old, mice were anesthetized with 200 mg/kg Tribromoethanol (Avertin) and transcardially perfused with TBS/Heparin and 4% paraformaldehyde (PFA) at room temperature (RT). Harvested brains were post-fixed overnight in 4% PFA, cryoprotected in 30% sucrose, and the brain blocks were prepared with O.C.T. (TissueTek) to store at −80°C. 30 µm thick brain sections were obtained through cryosectioning using a Leica CM3050S (Leica, Germany) vibratome and stored in a mixture of TBS and glycerol at −20°C for further free-float antibody staining procedures.

##### Immunohistochemistry on mouse brain sections

Mice used for immunohistochemistry were anesthetized with 200 mg/kg Avertin and perfused with TBS/Heparin and 4% PFA. Brains were collected and post-fixed in 4% PFA overnight, cryoprotected in 30% sucrose, frozen in a solution containing 2 parts 30% sucrose and 1-part O.C.T. (TissueTek), and stored at −80°C. Floating coronal tissue sections of 30 μm, 40 μm or 100 μm thickness were collected and stored in a 1:1 mixture of TBS/glycerol at −20°C. For immunostaining, sections were washed in 1x TBS containing 0.2% Triton X-100 (TBST), blocked in 10% NGS diluted in TBST, and incubated in primary antibody for 2-3 nights at 4°C with gentle shaking. Primary antibodies used were anti-LRRK2 (Rabbit, 1:500; ab133474, Abcam, RRID:AB_2713963), HA (Rat, 1:500; 11867423001, Roche, RRID:AB_390918), phospho-ERM (Rabbit, 1:500; 3141, Cell Signaling, RRID:AB_330232), Sox9 (Rabbit, 1:500; AB5535, Millipore Sigma, RRID:AB_2239761), GFAP (Rabbit, 1:500; Z0334, Agilent DAKO, RRID:AB_177521), VGluT1 (Guinea pig, 1:2000; 135304, Synaptic Systems, RRID: AB_887878), PSD95 (Rabbit, 1:300; 51-6900, Innovative Research, RRID: AB_87705), VGAT (Guinea pig, 1:1000; 131004, Synaptic Systems, RRID: AB_887873), Gephyrin (Rabbit, 1:1000; 147011, Synaptic Systems, RRID:AB_887717), S100β (Mouse IgG1, 1:200; S2532, Millipore Sigma, RRID: AB_477499), MAP2 (Mouse IgG1, 1:200; 13-1500, Thermo Fisher Scientific, RRID:AB_2533001), and Iba1 (Chicken, 1:500; 234009, Synaptic Systems, RRID: AB_2891282). For lambda protein phosphatase (λ-PPase) treatment, brain sections were divided into two groups: λ-PPase treatment group and a control group. Brain sections in the treatment group were incubated with λ-PPase at 30°C for 2 hours, while brain sections in the control group were incubated under identical conditions without the enzyme. All sections were blocked in 5% normal goat serum and 0.3% Triton X-100 for 1 hour at room temperature, followed by overnight incubation with primary antibodies at 4°C. Following the primary incubation, sections were washed in TBST, incubated in Alexa Fluor conjugated secondary antibodies diluted 1:200 (Life Technologies) for 2-3 hours at room temperature, washed with TBST, and mounted onto glass slides using a homemade mounting media (90% Glycerol, 20 mM Tris pH 8.0, 0.5% n-Propyl gallate) and sealed with nail polish. Images were acquired with an Olympus FV 3000 microscope.

##### Immunohistochemistry on human brain sections

Floating human frontal cortex sections of 40 μm thickness were obtained from Banner Sun Health Research Institute in Sun City, Arizona (4 control and 3 LRRK2 G2019S mutation carrier subjects). None of the control subjects had a history of dementia or neurological or psychiatric disorders at the time of death (See Supplemental Table 1). Informed and written consent was obtained from the donors. For immunostaining, sections were washed in 1x TBS containing 0.3% Triton X-100 (TBST), blocked in 3% NGS diluted in TBST, and incubated in primary antibody 2-3 nights at 4°C with shaking. Primary antibodies used were GFAP (chicken, 1:250; AB5541, Millipore Sigma, RRID:AB_177521), phospho-ERM (Rabbit, 1:250; 3726, Cell Signaling, RRID: AB_10560513), S100β (Mouse IgG1, 1:200; S2532, Millipore Sigma, RRID: AB_477499), and MAP2 (Mouse IgG1, 1:200; 13-1500, Thermo Fisher Scientific, RRID:AB_2533001). Following primary incubation, sections were washed in TBST, incubated in Alexa Fluor conjugated secondary antibodies diluted 1:200 (Life Technologies) for 2-3 hours at room temperature, washed with TBST, and mounted onto glass slides using a homemade mounting media (90% Glycerol, 20 mM Tris pH 8.0, 0.5% n-Propyl gallate) and sealed with nail polish.

##### Mouse astrocyte territory volume analysis

To assess the territory volume of individual astrocytes in the mouse cortex, 100 μm-thick floating sections containing anterior cingulate cortex (ACC) and primary motor cortex (MOp) astrocytes labeled sparsely via PALE with mCherry-CAAX were collected. High-magnification images containing an entire astrocyte (50-60 μm z-stack) were acquired with an Olympus FV 3000 microscope with the 60x objective. Criteria for data inclusion required that the entirety of the astrocyte could be captured within a single brain section and that the astrocyte was in L2-3 of the ACC or MOp. Astrocytes in which the entire astrocyte could not be captured within the section or was in other layers or outside of the ACC or MOp were not imaged. Imaged astrocytes were analyzed using Imaris Bitplane software (RRID: SCR_007370) as described previously[36]. Briefly, surface reconstructions were generated, and the Imaris Xtensions “Visualize Surface Spots” and “Convex Hull” were used to create an additional surface render representing the territory of the astrocyte. The volume of each territory was recorded, and astrocyte territory sizes from biological replicates were analyzed across experimental conditions using a nested two-way ANOVA followed by the Bonferroni posthoc test. For 3D Sholl analysis of individual PALE astrocytes, we first loaded images onto Imaris and then created a surface. After generating the surface of astrocytes, we created filaments using ‘Add new filament (leaf icon)’. For the quantification of complexity, we clicked on the gear tool on Imaris to display Sholl intersections. The number of animals and cells/animals analyzed are specified in the figure legend for each experiment.

##### Mouse synapse imaging and analysis

Staining, image acquisition, and analysis of synaptic density were performed as described previously[96]. Synaptic staining was performed in coronal sections (30 μm thick) containing the ACC and MOp from WT and LRRK2 G2019S^ki/ki^ mice. To label pre and postsynaptic proteins, the following antibody combinations were used: VGluT1 and PSD95 (excitatory, intracortical), VGAT, and GEPHYRIN (inhibitory). High magnification 60x objective plus 1.64x optical zoom z-stack images containing 15 optical sections spaced 0.34 μm apart were obtained using an Olympus FV 3000 inverted confocal microscope. Each z-stack was converted into 5 maximum projection images (MPI), each corresponding to a 1 μm section of z plane, using FIJI. Synapses were identified by the colocalization of pre and postsynaptic puncta. The number of co-localized synaptic puncta of excitatory intracortical (VGluT1-PSD95) and inhibitory (VGAT-GEPHYRIN) synapses were obtained using the FIJI plugin Puncta Analyzer[46]. 15 MPIs were analyzed for each mouse (5 MPI/tissue section, 3 tissue sections/mouse). Between 4 and 5, age and sex-matched mice/genotype/condition were analyzed for each synaptic staining combination, as indicated in the figure legends. All animals appeared healthy at the time of collection. No data were excluded.

##### Cell counting, imaging, and analysis

Tile scan images containing the anterior cingulated cortex (ACC) and primary motor cortex (MOp) from P21 WT and LRRK2 G2019S^ki/ki^; Aldh1l1-eGFP mice were acquired on a confocal Leica SP8 STED microscope using the galvo scanner and 20x objective. For ALDH1L1-EGFP and SOX9 double positive cell counting in the ACC and MOp, the cells labeled with ALDH1L1-EGFP and SOX9 were counted by hand using the cell counter tool in FIJI. 2-4 sections per brain from 3 sex and age-matched mice/genotype were analyzed.

#### Biochemical assays

##### mRNA extraction and cDNA preparation

Cells stored in TRIzol (15596026, Invitrogen) were thawed and resuspended in 1 mL of TRIzol. After adding 200 μL of chloroform, the samples were centrifuged at 12,000 g for 15 minutes at 4°C. The aqueous phase was collected, and RNA was precipitated with GlycoBlue Coprecipitant (AM9515, Invitrogen) and isopropanol. The RNA pellet was washed with 75% ethanol, air-dried, and resuspended in 40 μL of nuclease-free water. The RNA was then isolated using the Zymo Research RNA Clean & Concentrator-5 Kit (R1014, Zymo), quantified with the Qubit RNA HS Assay Kit (Q32852, Invitrogen), and diluted to equalize concentrations. Finally, cDNA libraries were generated using qScript cDNA SuperMix (101414-102, VWR) with a temperature profile of 25°C for 5 minutes, 42°C for 30 minutes, and 85°C for 5 minutes, and the resulting cDNA was diluted threefold and stored at −80°C.

##### Real-time qPCR

cDNA samples were plated on a 96-well qPCR plate and incubated with Fast SYBR Green Master Mix (4385616, Applied Biosystems), nuclease-free water, and the forward and reverse primers of interest at a ratio of 5 μl SYBR: 3 μl water: 0.5 μl forward primer: 0.5 μl reverse primer: 1 μl sample. Each sample was plated two to four times to ensure technical replicates. A control sample (water with primers and Master Mix) served as a negative control. Cycle threshold values were collected for each well and normalized to GAPDH as a housekeeping gene. The sequences of forward (F) and reverse (R) primers used (5′→ 3′) are: Atg7: (F) 5′- GTTCGCCCCCTTTAATAGTGC -3′ and (R) 5′- TGAACTCCAACGTCAAGCGG -3′

##### Protein extraction and Western blotting

Protein was extracted from cultured rat astrocytes using membrane solubilization buffer (25 mM Tris pH 7.4, 150 mM NaCl, 1 mM CaCl_2_, 1 mM MgCl_2_, 0.5% NP-40, and protease inhibitors). Cultured astrocytes were washed twice with ice-cold TBS containing 1 mM CaCl_2_ and 1 mM MgCl_2_ and incubated on ice in membrane solubilization buffer for 20 minutes with occasional agitation. Cell lysates were collected, vortexed briefly, and centrifuged at 4°C at high speed for 10 minutes to pellet non-solubilized material. The supernatant was collected and stored at −80°C.

Pierce BCA Protein Assay Kit (Thermo Fisher) was used to determine protein concentration, and lysates were mixed with 4x Pierce™ Reducing Sample Buffer (Thermo Scientific) and incubated at 45°C for 45 minutes to denature proteins. 7-10 μg (cultured astrocyte lysates) of protein was loaded into Bolt™ 4–12% Bus-Tris Plus gels (Thermo Scientific) and run at 150 V for 1 hour. Proteins were transferred at 100 V to PVDF membrane (Millipore) for 1.5 hours, blocked in 5% BSA in TBST (137 mM NaCl, 2.68 mM KCl, 24.7 mM Tris-Base, 0.1% (w/v) Tween 20) for 1 hour and incubated in primary antibodies overnight at 4°C. Primary antibodies used were: anti-LRRK2 (Rabbit, 1:500; ab133474, abcam, RRID:AB_2713963), GAPDH (mouse, 1:5000; ab8245, abcam, RRID:AB_2107448), β-actin (mouse, 1:5000; A5441, Millipore Sigma, RRID:AB_476744). The next day, membranes were washed with TBST, incubated in HRP secondary antibodies (Thermo Fisher Scientific) for 2 hours, washed in TBST, and imaged on a Biorad Gel Doc imaging system. Protein levels were quantified using FIJI.

##### Coimmunoprecipitation

Co-immunoprecipitation assays were performed in HEK293T cells. Cells were transfected 36 hours prior to lysis with Ezrin/Atg7 cDNA using X-tremeGENE HP (Roche) and grown to 85-90% confluency. At 36 hours post-transfection, cells were rinsed with 1x PBS and collected for lysis. Cellular lysis and protein extraction was conducted via brief vortexing in chilled membrane solubilization buffer (25mM HEPES, 150mM KCl, 1.5mM MgCl2, 0.5% NP40, 10% Glycerol) in the presence of both protease (cOmplete, Roche) and phosphatase (PhosSTOP, Roche) inhibitors. Protein concentrations of cell lysates were equalized using the Pierce™ BSA Protein Assay Kit (Thermo Fisher) and CLARIOstar Plus Plate Reader (BMG Labtech). Equalized cell lysates were incubated, while rotating, with Pierce™ Anti-c-Myc Magnetic Beads (Thermo Fisher) for 6 hours at 4°C. Following this incubation, the beads were washed with chilled lysis buffer. Protein samples were eluted from beads through the addition of 2x Bromophenol Blue-free Laemilli Sample Buffer and heating to 95°C for 5 minutes. Eluted protein samples were subjected to western blot analysis and quantification using a SimpleWestern Jess (ProteinSimple) automated immunoassay system with a 12-230kDa Fluorescence Separation module and associated manufacturer’s protocol. Primary antibodies used for protein detection were: Anti-Myc (Rabbit, 1:40, Cell Signaling Technologies, mAb#2278) and Anti-HA (Rat, 1:20, Roche, 11867423001). Signal detection was achieved using the following fluorescently conjugated secondary antibodies at a dilution of 1:100: IRDye® 680RD Goat anti-Rabbit IgG (LI-COR, 926-68071) and IRDye® 680RD Goat anti-Rat IgG (LI-COR, 926-68076). Signal quantification was conducted with the Simple Western Compass software (https://www.bio-techne.com/resources/instrument-software-download-center/compass-software-simple-western, RRID: SCR_022930), utilizing quantitative electropherograms of detected signals. Ezrin co-immunoprecipitation signal intensity was background-subtracted and normalized to Ezrin protein load and Atg7 IP levels for statistical analysis. Statistical analysis was performed in GraphPad Prism 9, using One-way ANOVA with Tukey’s multiple comparisons test and alpha threshold of 0.05.

##### Protein interaction modeling

Full-length mouse Ezrin (UniProt ID: Q4KML7) and Atg7 (UniProt ID: Q9D906) were used for structure prediction in Alphafold 2.0 Multimer. The structure of Ezrin in the open conformation was generated by segmenting independently predicted Ezrin N- and C-termini and threading them onto an open ERM hinge structure using the PyMOL2 molecular visualization software. To model the Ezrin:Atg7 interaction and associated conformational changes, the structures of Ezrin and the Atg7 homodimer were predicted and docked using Alphafold 2.0 Multimer. Conformational changes in the Ezrin:Atg7 protein complex were assessed using PyMOL2 and CABS-flex 2.0[97]. For all predicted structure models, the highest-confidence structures, calculated via the predicted local distance difference test (pLDDT), were subjected to energy-minimization via AMBER relaxation. The resulting minimized structures were imported into PyMOL2 for representative model production.

#### Proteomic analysis

##### *In vivo* BioID protein purification

*In vivo,* BioID experiments were performed as described in Takano et al., 2020 with modifications[98]. Genotype-matched animals (wild-type C57BL6 or LRRK2 G2019S^ki/ki^) were bred to produce single-genotype litters. For each genotype (WT or G2019S), 6 pups were injected with AAVs carrying Astro-Ezrin-BioID (PHP.eB.GfaABC1D-Ezrin WT-BioID2-HA) or Astro-CYTO-BioID (PHP.eB.GfaABC1D-BioID2-HA). 2 genotype-matched cortices were pooled at the time of protein isolation, yielding 3 independent replicates per BioID construct. 12 total samples (3 per genotype per construct) were subjected to LC-MS/MS and downstream analysis. WT and LRRK2 G2019S^ki/ki^ P0 – P2 mouse pups were anesthetized by hypothermia, and 1µl of each concentrated AAV-BioID vector was injected bilaterally into the cortex using a Hamilton syringe. Pups were monitored until they recovered on a heating pad. At P18, P19, and P20, biotin was subcutaneously injected at 24 mg/kg to increase the biotinylation efficiency. At P21, the cerebral cortices were removed and stored at -80° C. For the protein purification, each cortex was lysed in a buffer containing 50 mM Tris/HCl, pH 7.5; 150 mM NaCl; 1 mM EDTA; protease inhibitor mixture (Roche); and phosphatase inhibitor mixture (PhosSTOP, Roche). Next, an equal volume of buffer containing 50 mM Tris/HCl, pH 7.5; 150 mM NaCl; 1 mM EDTA; 0.4 % SDS; 2 % TritonX-100; 2 % deoxycholate; protease inhibitor mixture; and phosphatase inhibitor mixture was added to the samples, following by sonication and centrifugation at 15,000 g for 10 min. The remaining supernatant was ultracentrifuged at 100,000g for 30min at 4° C. Finally, SDS detergent was added to the samples and heated at 45 ° C for 45 min. After cooling on ice, each sample was incubated with High-Capacity Streptavidin Agarose beads (ThermoFisher) at 4° C overnight. Following incubation, the beads were serially washed: 1) twice with a solution containing 2% SDS; 2) twice with a buffer 1% TritonX-100, 1% deoxycholate, 25 mM LiCl; 3) twice with 1M NaCl, and finally, five times with 50 mM ammonium bicarbonate. The biotinylated proteins attached to the agarose beads were eluted in a buffer of 125 mM Tris/HCl, pH6.8; 4 % SDS; 0.2 % β-mercaptoethanol; 20 % glycerol; 3 mM biotin at 60°C for 15min.

##### Sample Preparation

The Duke Proteomics and Metabolomics Shared Resource (DPMSR) received 12 samples (3 of each WT CYT, GS CYT, WT EZR, and GS EZR) which were kept at -80C until processing. Samples were spiked with undigested bovine casein at a total of either 1 or 2 pmol as an internal quality control standard. Next, they were reduced for 15 min at 80C, alkylated with 20 mM iodoacetamide for 30 min at room temperature, then supplemented with a final concentration of 1.2% phosphoric acid and 636 μl of S-Trap (Protifi) binding buffer (90% MeOH/100mM TEAB). Proteins were trapped on the S-Trap micro cartridge, digested using 20 ng/μL sequencing grade trypsin (Promega) for 1 hr at 47C, and eluted using 50 mM TEAB, followed by 0.2% FA, and lastly using 50% ACN/0.2% FA. All samples were then lyophilized to dryness. Samples were resuspended in 12 μl of 1% TFA/2% acetonitrile with 12.5 fmol/μl of yeast ADH. A study pool QC (SPQC) was created by combining equal volumes of each sample.

##### LC-MS/MS Analysis

Quantitative LC/MS/MS was performed on 3 μL of each sample, using an MClass UPLC system (Waters Corp) coupled to a Thermo Orbitrap Fusion Lumos high resolution accurate mass tandem mass spectrometer (Thermo) equipped with a FAIMSPro device via a nanoelectrospray ionization source. Briefly, the sample was first trapped on a Symmetry C18 20 mm × 180 μm trapping column (5 μl/min at 99.9/0.1 v/v water/acetonitrile), after which the analytical separation was performed using a 1.8 μm Acquity HSS T3 C18 75 μm × 250 mm column (Waters Corp.) with a 90-min linear gradient of 5 to 30% acetonitrile with 0.1% formic acid at a flow rate of 400 nanoliters/minute (nL/min) with a column temperature of 55C. Data collection on the Fusion Lumos mass spectrometer was performed for three difference compensation voltages (-40v, -60v, -80v). Within each CV, a data-dependent acquisition (DDA) mode of acquisition with a r=120,000 (@ m/z 200) full MS scan from m/z 375 – 1500 with a target AGC value of 4e5 ions was performed. MS/MS scans with HCD settings of 30% were acquired in the linear ion trap in “rapid” mode with a target AGC value of 1e4 and max fill time of 35 ms. The total cycle time for each CV was 0.66s, with total cycle times of 2 sec between like full MS scans. A 20s dynamic exclusion was employed to increase the depth of coverage. The total analysis cycle time for each sample injection was approximately 2 hours.

##### Quantitative Data Analysis

Following 15 total UPLC-MS/MS analyses (excluding conditioning runs, but including 3 replicate SPQC samples), data were imported into Proteome Discoverer 3.0 (Thermo Scientific Inc.) and individual LCMS data files were aligned based on the accurate mass and retention time of detected precursor ions (“features”) using Minora Feature Detector algorithm in Proteome Discoverer. Relative peptide abundance was measured based on peak intensities of selected ion chromatograms of the aligned features across all runs. The MS/MS data was searched against the SwissProt *M. musculus* database, a common contaminant/spiked protein database (bovine albumin, bovine casein, yeast ADH, etc.), and an equal number of reversed-sequence “decoys” for false discovery rate determination. Sequest with INFERYS was utilized to produce fragment ion spectra and to perform the database searches. Database search parameters included fixed modification on Cys (carbamidomethyl) and variable modification on Met (oxidation). Search tolerances were 2ppm precursor and 0.8Da product ion with full trypsin enzyme rules. Peptide Validator and Protein FDR Validator nodes in Proteome Discoverer were used to annotate the data at a maximum 1% protein false discovery rate based on q-value calculations. Note that peptide homology was addressed using razor rules in which a peptide matched to multiple different proteins was exclusively assigned to the protein has more identified peptides. Protein homology was addressed by grouping proteins that had the same set of peptides to account for their identification. A master protein within a group was assigned based on % coverage.

Prior to normalization, a filter was applied such that a peptide was removed if it was not measured at least twice across all samples and in at least 50% of the replicates in any one single group. After that filter, samples were total intensity normalized (total intensity of all peptides for a sample are summed and then normalized across all samples). Next, the following imputation strategy is applied to missing values. If less than half of the values are missing in a biological group, values are imputed with an intensity derived from a normal distribution of all values defined by measured values within the same intensity range (20 bins). If greater than half values are missing for a peptide in a group and a peptide intensity is > 5e6, then it was concluded that the peptide was misaligned and its measured intensity is set to 0. All remaining missing values are imputed with the lowest 2% of all detected values. Peptide intensities were then subjected to a trimmed-mean normalization in which the top and bottom 10 percent of the signals were excluded, and the average of the remaining values was used to normalize across all samples. Lastly, all peptides belonging to the same protein were then summed into a single intensity. These normalized protein level intensities are what was used for the remainder of the analysis.

To assess technical reproducibility, we calculated the % coefficient of variation (%CV) for each protein across 3 injections of an SPQC pool that were interspersed throughout the study (Table S4). The mean %CV of the SPQC pools was 11.7%, which is well within our expected analytical tolerances. To assess biological + technical variability, %CVs were measured for each protein across the individual groups which averaged 19.8%, which is also within our normal tolerances and indicated a reproducible IP and sample processing.

As an initial statistical analysis, we calculated fold-changes between various sample groups based on the protein expression values and calculated two-tailed heteroscedastic t-test on log2-transformed data for this comparison. Those fold changes and p-values are presented for all proteins in Tables S5, S7, S9, and S10. False Discovery Rate (FDR) correction was performed using the Benjamini-Hochberg procedure, and these values are reported (“p.adjusted”) in Tables S5, S7, S9, and S10. To optimize for discovery, we opted to use the less stringent parameter of unadjusted p-value for downstream analyses. Within supplemental tables, we have labeled proteins significantly more abundant (“Up”, fold change >1.5 and unadjusted p-value <0.05), or less abundant (“Down”, fold change >-1.5 and unadjusted p-value <0.05) in a particular genotype or BioID sample group. These annotated proteins were used for downstream analyses, including protein interaction networks using Cytoscape (v.3.9.1) and Gene Ontology (GO) enrichment analysis using the ClusterProfiler package for R, with all M. musculus genes as the reference background.

##### Mouse brain slice electrophysiology

For whole-cell patch-clamp recordings, 3-4 mice of each genotype and condition were used for miniature excitatory postsynaptic current (mEPSC) and miniature inhibitory postsynaptic current (mIPSC) measurements. WT and *LRRK2* G2019S^ki/ki^ mice of both sexes were anesthetized with 200 mg/kg tribromoethanol (avertin) and decapitated. After decapitation, the brains were immersed in ice-cold artificial cerebrospinal fluid (aCSF, in mM): 125 NaCl, 2.5 KCl, 3 mM MgCl_2_, 0.1 mM CaCl_2_, 10 glucose, 25 NaHCO_3_, 1.25 NaHPO_4_, 0.4 L-ascorbic acid, and 2 Na-pyruvate, pH 7.3-7.4 (310 mOsmol). 350 μm thick coronal slices containing the ACC were obtained using a vibrating tissue slicer (Leica VT1200; Leica Biosystems). Slices were immediately transferred to standard aCSF (33°C, continuously bubbled with 95% O_2_ – 5% CO_2_) containing the same as the low-calcium aCSF but with 1 mM MgCl_2_ and 1-2 mM CaCl_2_. After 30-minute incubation at 33°C, slices were transferred to a holding chamber with the same extracellular buffer at room temperature (RT: ∼25°C). Brain slices were visualized by an upright microscope (BX61WI, Olympus) through a 40x water-immersion objective equipped with infrared-differential interference contrast optics in combination with a digital camera (ODA-IR2000WCTRL). Patch-clamp recordings were performed by using an EPC 10 patch-clamp amplifier, controlled by Patchmaster Software (HEKA). Data were acquired at a sampling rate of 50 kHz and low-pass filtered at 6 kHz.

To measure mEPSCs, the internal solution contained the following (in mM): 125 K-gluconate, 10 NaCl, 10 HEPES, 0.2 EGTA, 4.5 MgATP, 0.3 NaGTP, and 10 Na-phosphocreatine, pH adjusted to 7.2 – 7.4 with KOH and osmolality set to ∼ 300 mOsmol. mEPSCs were measured in the aCSF bath solution containing 1 µM tetrodotoxin and 50 µM Picrotoxin at -70 mV in voltage-clamp mode. To measure mIPSCs, the internal solution contained the following (in mM): 77 K-gluconate, 77 KCl, 10 HEPES, 0.2 EGTA, 4.5 MgATP, 0.3 NaGTP, and 10 Na-phosphocreatine, pH adjusted to 7.2 – 7.4 with KOH and osmolality set to ∼ 300 mOsmol. mIPSCs were measured in the aCSF bath solution containing 1 µM tetrodotoxin and 10 µM 6-cyano-7-nitroquinoxaline-2,3-dione (CNQX), and 50 µM D-2-amino-5-phosphonopentanoate (D-AP5) at -70 mV in voltage-clamp mode. mEPSCs and mIPSCs recorded at -70 mV were detected using Minhee Analysis software (https://github.com/parkgilbong/Minhee_Analysis_Pack). To analyze the frequency, events were counted over 5 minutes of recording. To obtain the average events for each cell, at least 100 non-overlapping events were detected and averaged. The peak amplitude of the average mEPSCs was measured relative to the baseline current.

##### Quantification and statistical analysis

All statistical analyses were performed in GraphPad Prism 9. Exact value of n, what n represents, and specific statistical tests for each experiment are indicated in the figure legend for each experiment. All data are represented as mean ± standard error of the mean, and data points are shown where applicable. Exact *P*-values are listed in the figure for each experiment. Where indicated, the unpaired two-tailed t-tests were run using Welch’s correction, and a correction for multiple comparisons was applied using Hom-Sidak method with an alpha threshold of 0.05 for adjusted *P*-value. A Geisser-Greenhouse correction was used for both One-way and Two-way ANOVA analyses. For Sholl analysis, statistical analyses were conducted using a custom code in R (https://github.com/Eroglu-Lab/In-Vitro-Sholl). A mixed-effect model with multiple comparisons made using the Tukey post-test was used for Sholl analysis to account for the variability per experiment as a random effect to evaluate differences between the conditions. Sample sizes were determined based on previous experience for each experiment to yield high power to detect specific effects. No statistical methods were used to predetermine the sample size. All experimental animals that appeared healthy at the time of tissue collection were processed for data collection.

## ACKNOWLEDGMENTS

We thank Drs. Amy Lee, Zhiyong Liu, Nicole Calakos, Katja Brose, and Andrew Singleton for critical feedback about the manuscript. We thank Adam Okinaga and Donna Porter for their excellent technical help. This work was supported by a Foerster Bernstein Women In STEM Postdoctoral Fellowship to SW, and a Chan Zuckerberg Initiative, Neurodegeneration Challenge Network Collaborative grant (DAF2018-191999 and DAF2021-237435) to CE and ARLS, and by the N.I.H. grants R35-NS122140 to A.R.L.S. and R01-NS102237 to CE, and by the Michael J. Fox Foundation for Parkinson’s Research (MJFF) Astrocyte Biology in PD pilot grant (MJFF 17914 to CE and ARLS), and by the joint efforts of MJFF and the Aligning Science Across Parkinson’s (ASAP) initiative. MJFF administers the grant [ASAP-020607 to CE and SHS] on behalf of ASAP and the Michael J Fox Foundation. Dr. Cagla Eroglu is an HHMI Investigator. We are grateful to the Banner Sun Health Research Institute Brain and Body Donation Program of Sun City, Arizona for the provision of human biological materials. The Brain and Body Donation Program has been supported by the National Institute of Neurological Disorders and Stroke (U24 NS072026 National Brain and Tissue Resource for Parkinson’s Disease and Related Disorders), the National Institute on Aging (P30AG19610 and P30AG072980, Arizona Alzheimer’s Disease Center), the Arizona Biomedical Research Commission contracts 4001, 0011, 05-901 and 1001 to the Arizona Parkinson’s Disease Consortium and the Michael J. Fox Foundation for Parkinson’s Research.

## IP RIGHTS NOTICE

This article is subject to HHMI’s Open Access to Publications policy. HHMI lab heads have previously granted a nonexclusive CC BY 4.0 license to the public and a sublicensable license to HHMI in their research articles. Pursuant to those licenses, the author-accepted manuscript of this article can be made freely available under a CC BY 4.0 license immediately upon publication.

## AUTHOR CONTRIBUTIONS

Conceptualization, SW, ARLS, and CE; Methodology, SW, RB, GS, DSB, LV, KS, JM, CXT, TT, MPR, KD, NB, LB, SHS, and CE; Investigation, SW, RB, GS, and LV; Formal analysis, SW, RB, GS, and LV; Resources, CXT, MPR, and KS; Writing – original draft, SW and CE; Writing – Review & Editing, SW, RL, and CE; Funding Acquisition, SW, SHS, ARLS, and CE.

## Supplemental information

**Figure S1 (related to Figure 1).**
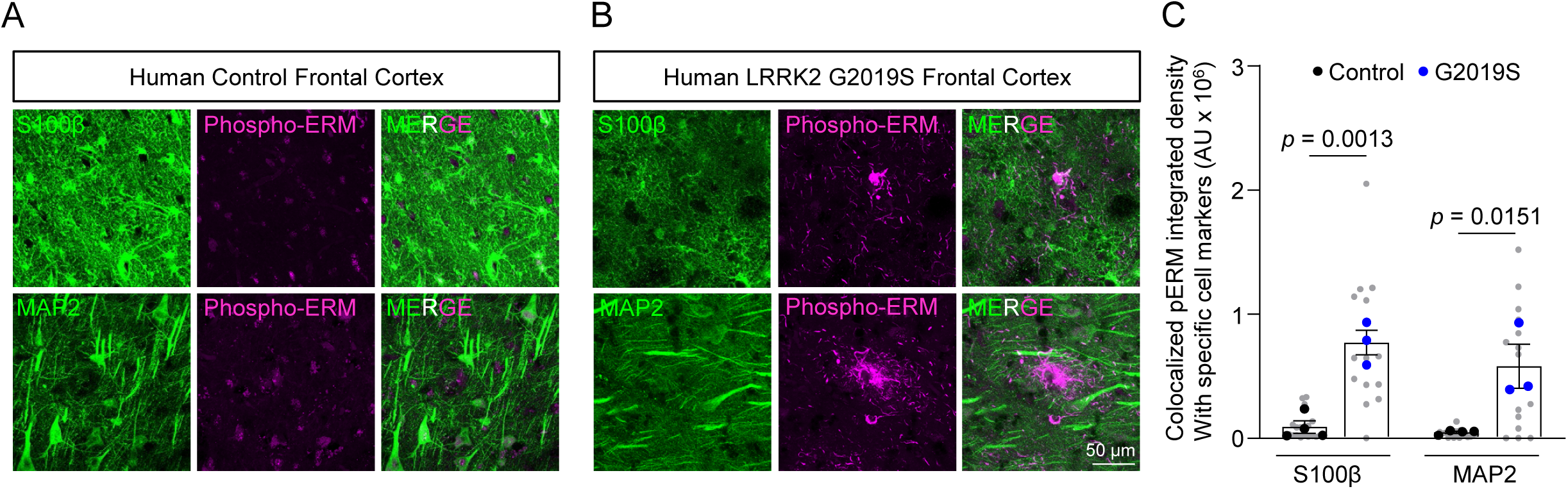
Human phospho-ERM is significantly increased and colocalizes with S100β and MAP2. Related to Figure 1. **(A-B)** Representative confocal images of phospho-ERM (purple) and S100β (green) or MAP2 (green) in the frontal cortex of human control subjects or human PD patients carrying LRRK2 G2019S mutation carriers at age >80 years old. Scale bar, 50 μm. **(C)** Quantification of phospho-ERM integrated density in (A-B), n = 4 (Human control, 3 males and 1 female), 3 (LRRK2 G2019S mutation carriers, 2 males and 1 female) subjects, nested t-test, unpaired two-tailed t-test. For colocalized phospho-ERM with S100β, t (5) = 6.541, *p* = 0.0013. For colocalized phospho-ERM with MAP2, t (5) = 3.625, *p* = 0.0151. Grey dots are the data acquired from each image. Black dots are the averaged data acquired from each control subject. Blue dots are the averaged data acquired from each LRRK2 G2019S mutation carrier.

**Figure S2 (related to Figure 1).**
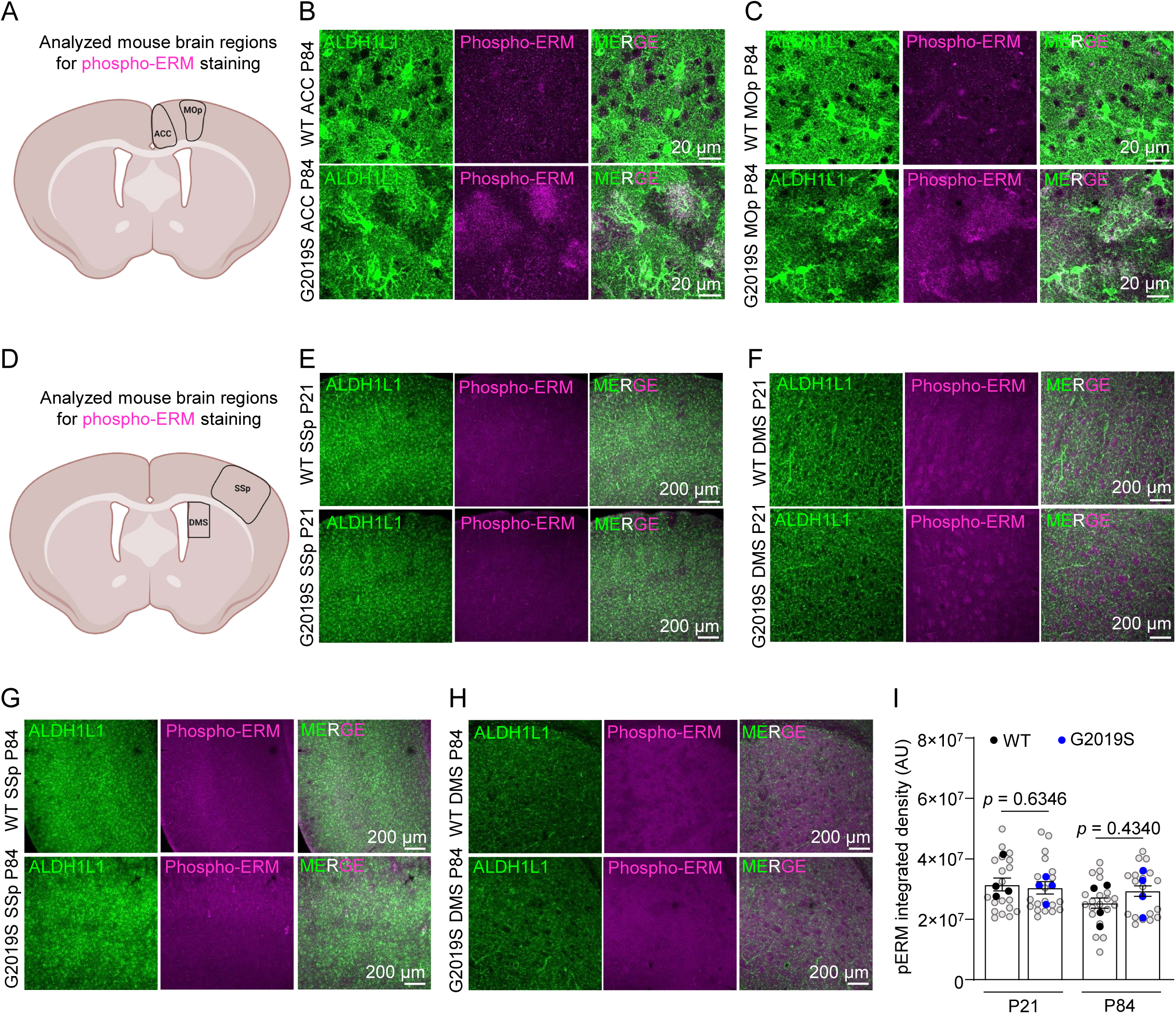
Phospho-ERM is upregulated in LRRK2 G2019S^ki/ki^ astrocytes in ACC and MOp but not SSp and DMS. Related to Figure 1. **(A)** Schematic representation of analyzed mouse brain regions for phospho-ERM staining, the ACC, and the MOp. **(B-C)** Representative confocal images of ERM phosphorylation (purple) in the ACC and MOp of WT or LRRK2 G2019S^ki/ki^ Aldh1L1-eGFP mice at P84. Scale bar, 20 μm. **(D)** Schematic representation of analyzed mouse brain regions for phospho-ERM staining, the somatosensory cortex (SSp), and the dorsal medial striatum (DMS). **(E-H)** Representative confocal images of ERM phosphorylation (purple) in the SSp and DMS of WT or LRRK2 G2019S^ki/ki^ Aldh1L1-eGFP mice at P21 (E-F) and at P84 (G-H). Scale bar, 200 μm. **(I)** Quantification of phospho-ERM integrated density in (E-H). For phospho-ERM integrated density at P21, nested t-test, unpaired two-tailed t-test. t (6) = 0.5005, *p* = 0.6346. n = 4 (WT, 2 males and 2 females), 4 (LRRK2 G2019S^ki/ki^, 2 males and 2 females) mice. For phospho-ERM integrated density at P84, nested t-test, unpaired two-tailed t-test. t (6) = 0.8383, *p* = 0.1123. n = 4 (WT, 2 males and 2 females), 4 (LRRK2 G2019S^ki/ki^, 2 males and 2 females) mice.

**Figure S3 (related to Figure 1).**
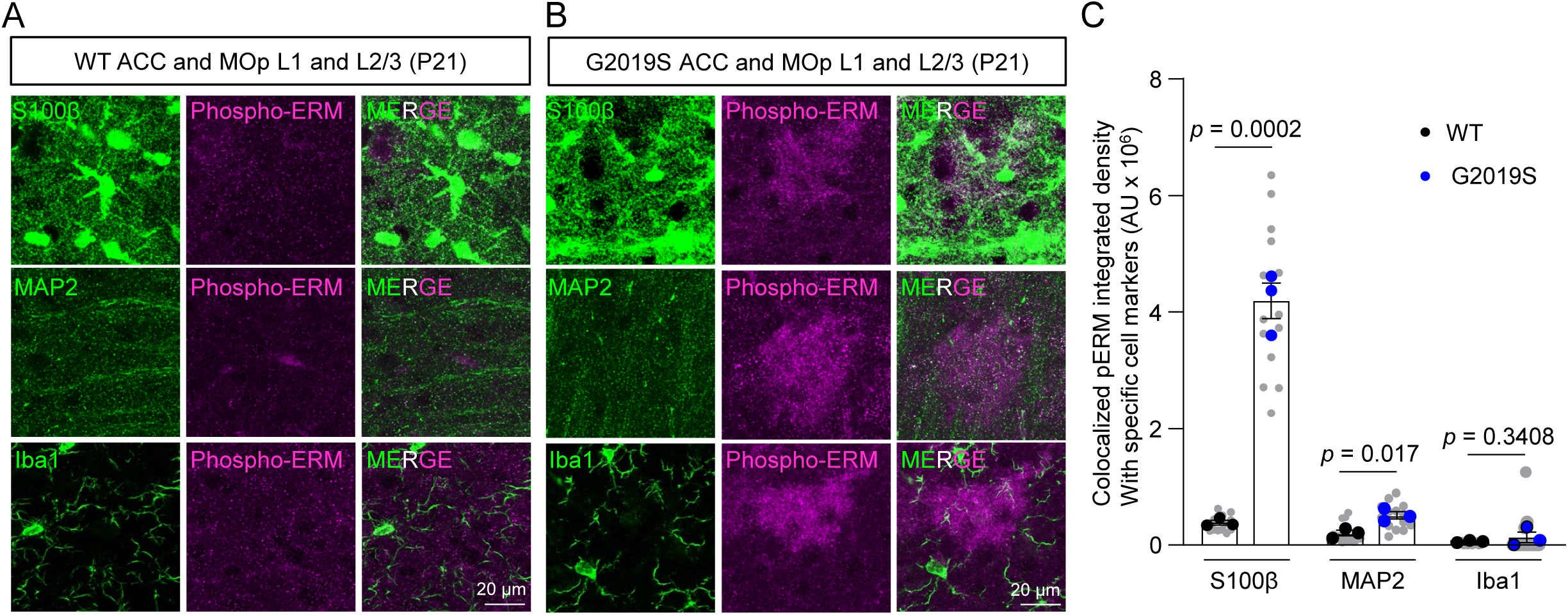
mouse phospho-ERM is significantly increased and colocalizes with S100β and MAP2. Related to Figure 1. **(A-B)** Representative confocal images of phospho-ERM (purple) and S100β (green), MAP2 (green), or Iba1 (green) from the ACC and MOp of WT or LRRK2 G2019S^ki/ki^ mice at P21. Scale bar, 20 μm. **(C)** Quantification of colocalized phospho-ERM integrated density with S100β, MAP2, and Iba1 in (D-E), n = 3 (WT, 1 male and 2 females), 3 (LRRK2 G2019S^ki/ki^, 1 male and 2 females) mice, nested t-test, unpaired two-tailed t-test. For colocalized phospho-ERM with S100β, t (4) = 12.37, *p* = 0.0002. For colocalized phospho-ERM with MAP2, t (4) = 3.939, *p* = 0.017. For colocalized phospho-ERM with Iba1, t (4) = 1.08, *p* = 0.3408. Grey dots are the data acquired from each image. Black dots are the averaged data acquired from each WT mouse. Blue dots are the averaged data acquired from each LRRK2 G2019S^ki/ki^ mouse.

**Figure S4 (related to Figure 1).**
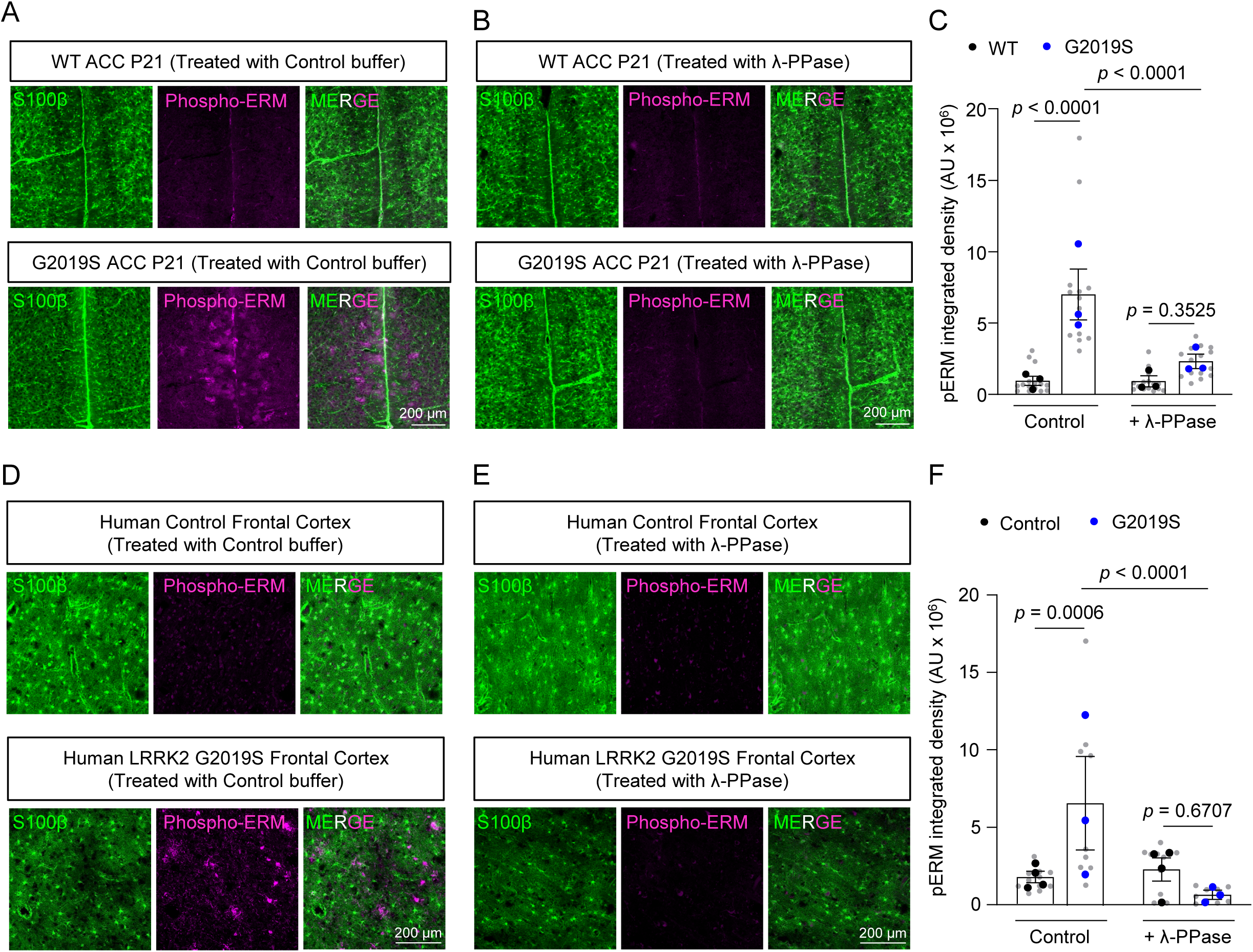
phospho-ERM is eliminated after Lambda protein phosphatase (λ-PPase) treatment. Related to Figure 1. **(A-B)** Representative confocal images of S100β (green) and phospho-ERM (purple) in the ACC of WT or LRRK2 G2019S^ki/ki^ mice at P21 with or without λ-PPase treatment. Scale bar, 200 μm. **(C)** Quantification of phospho-ERM integrated density in (A-B), n = 3 (WT, 1 male and 2 females), 3 (LRRK2 G2019S^ki/ki^, 1 male and 2 females) mice, Nested One-way ANOVA [F (3, 56) = 25.19, *p* < 0.0001], Bonferroni’s multiple comparisons test revealed a significant difference between WT mice and LRRK2 G2019S^ki/ki^ mice without λ-PPase treatment (*p* < 0.0001, 95% C.I. = [-8107799, -3924949]), and between LRRK2 G2019S^ki/ki^ mice with and without λ-PPase treatment (*p* < 0.0001, 95% C.I. = [2605448, 6788298]). alpha = 0.05. Grey dots are the data acquired from each image. Black dots are the averaged data acquired from each WT mouse. Blue dots are the averaged data acquired from each LRRK2 G2019S^ki/ki^ mouse. **(D-E)** Representative confocal images of S100β (green) and phospho-ERM (purple) in the frontal cortex of human control subjects or human PD patients carrying LRRK2 G2019S mutation carriers at age >80 years old with or without λ-PPase treatment. Scale bar, 200 μm. **(F)** Quantification of phospho-ERM integrated density in (D-E), n = 4 (Human control, 3 males and 1 female), 3 (LRRK2 G2019S mutation carriers, 2 males and 1 female) subjects, Nested One-way ANOVA [F (3, 38) = 9.296, *p* < 0.0001], Bonferroni’s multiple comparisons test revealed a significant difference between human control subjects and human PD patients carrying LRRK2 G2019S mutation carriers without λ-PPase treatment (*p* = 0.0006, 95% C.I. = [-7928018, - 1867234]), and between human PD patients carrying LRRK2 G2019S mutation carriers with and without λ-PPase treatment (*p* < 0.0001, 95% C.I. = [2768186, 9247437]). alpha = 0.05. Grey dots are the data acquired from each image. Black dots are the averaged data acquired from each control subject. Blue dots are the averaged data acquired from each LRRK2 G2019S mutation carrier.

**Figure S5 (related to Figure 1).**
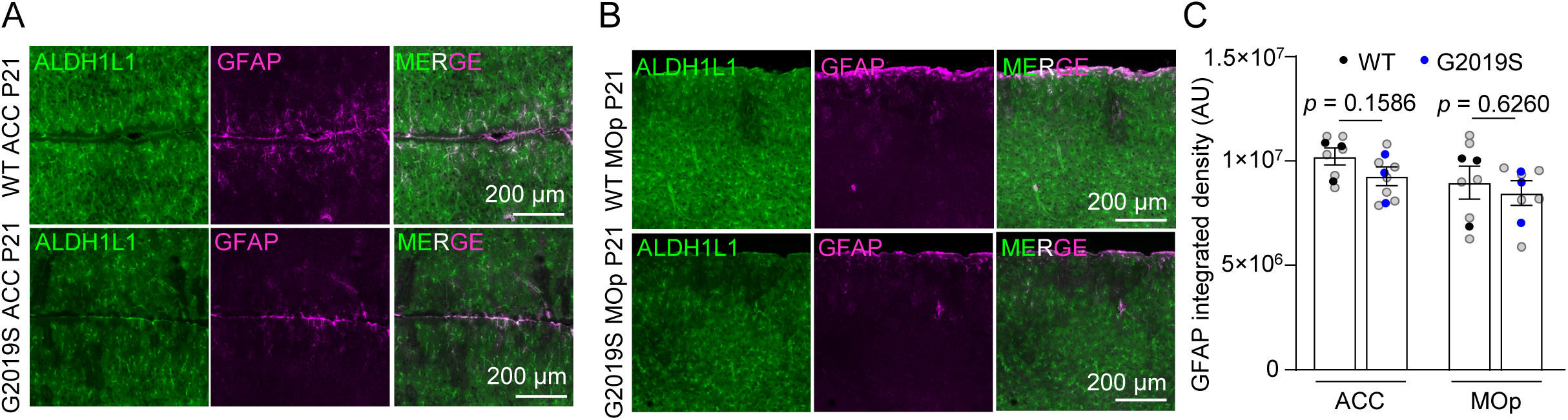
GFAP level is not altered in LRRK2 G2019S^ki/ki^ ACC and MOp. Related to Figure 1. **(A-B)** Representative confocal images of GFAP (purple) in the ACC and MOp of WT or LRRK2 G2019S^ki/ki^ Aldh1l1-eGFP mice at P21. Scale bar, 200 μm. **(C)** Quantification of GFAP integrated density in (A-B), n = 3 (WT, 2 males and 1 female), 3 (LRRK2 G2019S^ki/ki^, 2 males and 1 female) mice, For GFAP quantification in the ACC, nested t-test, unpaired two-tailed t-test. t (4) = 1.023, *p* = 0.364. For GFAP quantification in the MOp, nested t-test, unpaired two-tailed t-test. t (4) = 0.375, *p* = 0.7267. Grey dots are the data acquired from each image. Black dots are the averaged data acquired from each WT mouse. Blue dots are the averaged data acquired from each LRRK2 G2019S^ki/ki^ mouse.

**Figure S6 (related to Figure 2).**
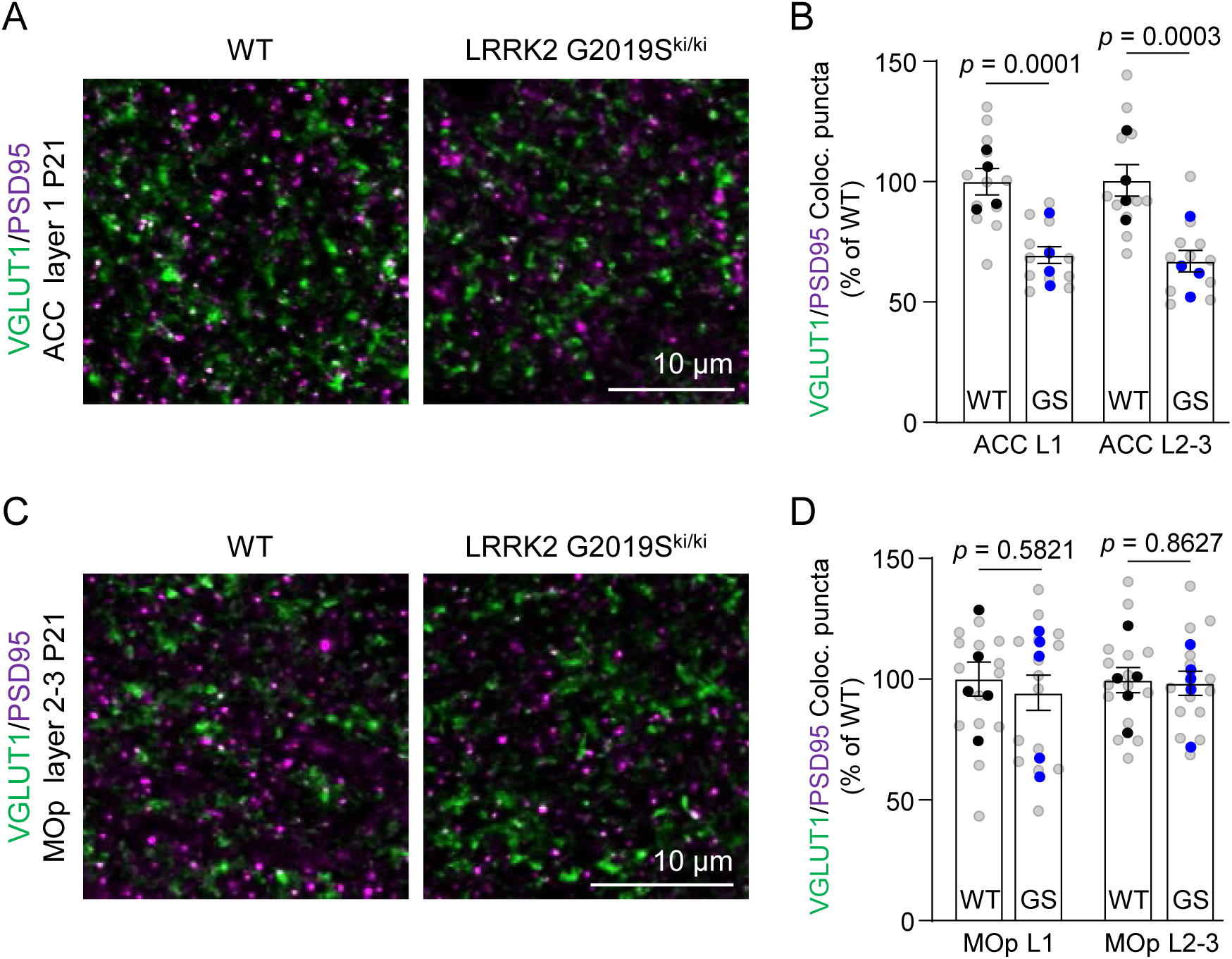
LRRK2 G2019S does not change excitatory synapse numbers in the primary motor cortex. Related to Figure 2. **(A)** Representative images from the ventral ACC of WT and LRRK2 G2019S^ki/ki^ mice that were stained with VGluT1 and PSD95 antibodies at P21. Scale bar, 10 μm. **(B)** Quantification of VGluT1-PSD95 co-localized puncta, normalized using the means of WT values in the ventral ACC. n = 4 (WT, 2 males and 2 females), 4 (LRRK2 G2019S^ki/ki^, 2 males and 2 females) mice. Nested One-way ANOVA [F (3, 44) = 12.98, *p* < 0.0001], Bonferroni’s multiple comparisons test revealed a significant difference between WT ACC L1 and LRRK2 G2019S^ki/ki^ ACC L1 (*p* = 0.0002, 95% C.I. = [13.71, 47.33]), and between WT ACC L2-3 and LRRK2 G2019S^ki/ki^ ACC L2-3 (*p* < 0.0001, 95% C.I. = [16.48, 50.10]). alpha = 0.05. **(C)** Representative images from the MOp of WT and LRRK2 G2019S^ki/ki^ mice that were stained with VGluT1 and PSD95 antibodies at P21. Scale bar, 10 μm. **(D)** Quantification of VGluT1-PSD95 co-localized puncta, normalized using the means of WT values in the MOp L1 and L2-3. n = 5 (WT), 5 (LRRK2 G2019S^ki/ki^) mice, 3 males and 2 females. Nested One-way ANOVA [F (3, 56) = 0.1837, *p* = 0.9070], Bonferroni’s multiple comparisons test revealed a significant difference between WT MOp L1 and LRRK2 G2019S^ki/ki^ MOp L1 (*p* > 0.9999, 95% C.I. = [-14.69, 25.97]), and between WT MOp L2-3 and LRRK2 G2019S^ki/ki^ MOp L2-3 (*p* > 0.9999, 95% C.I. = [-19.06, 21.61]). alpha = 0.05. Grey dots are the data acquired from each image. Black dots are the averaged data acquired from each WT mouse. Blue dots are the averaged data acquired from each LRRK2 G2019S^ki/ki^ mouse.

**Figure S7 (related to Figure 4).**
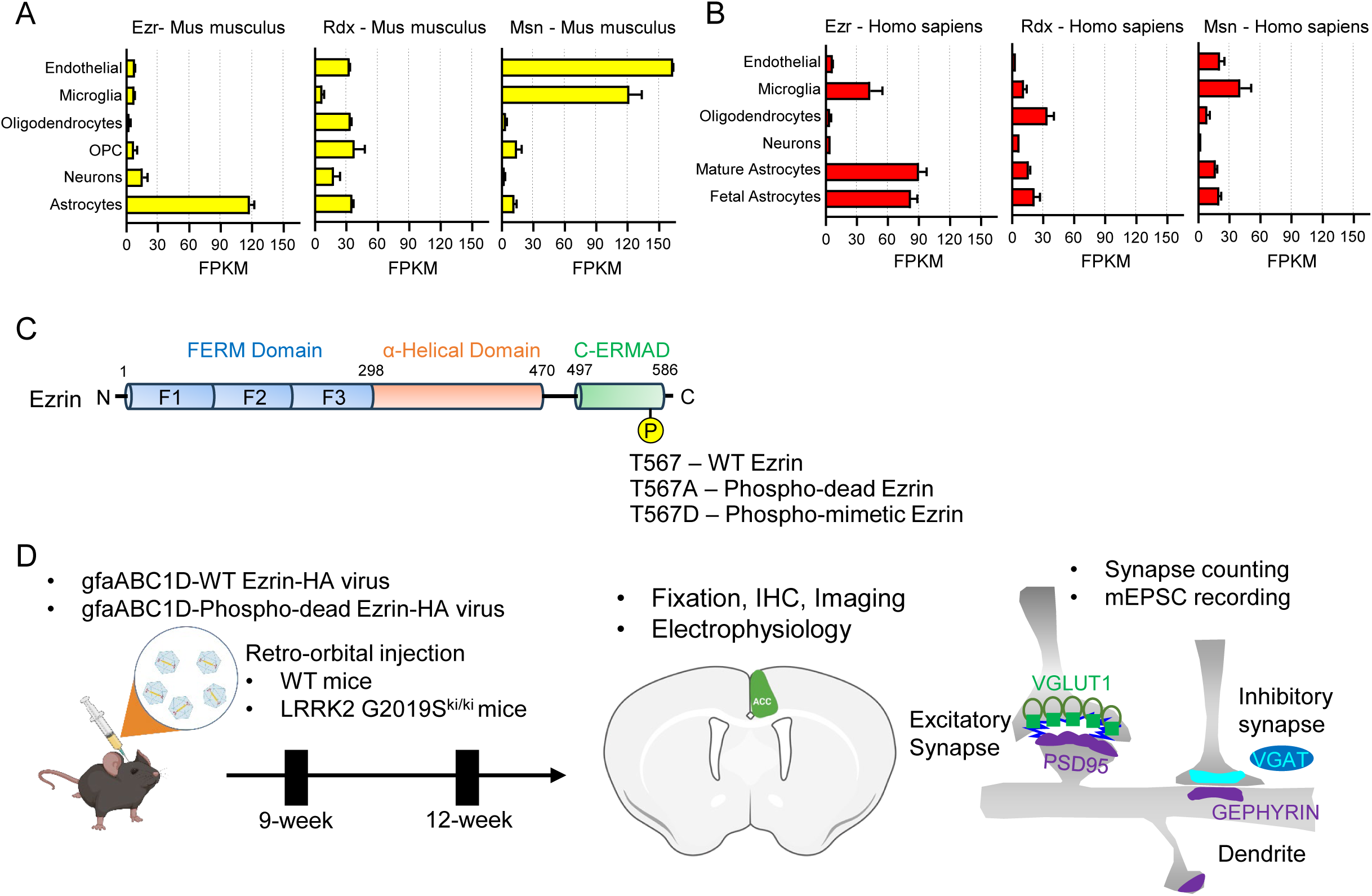
Overexpression of AAV-HA-tagged-WT Ezrin and AAV-HA-tagged-Phospho-dead Ezrin in WT and LRRK2 G2019S^ki/ki^ astrocytes. Related to Figure 4. **(A-B)** Bar plots of Ezr, Rdx, and Msn gene expression in various brain cell types in mice (yellow) or humans (red). These plots are generated using publicly available data at https://www.brainrnaseq.org **(C)** Schematic of domains within Ezrin. **(D)** Experiment workflow of AAV-HA-tagged-WT Ezrin and AAV-HA-tagged-Phospho-dead Ezrin in WT and LRRK2 G2019S^ki/ki^ astrocytes.

**Figure S8 (related to Figure 4).**
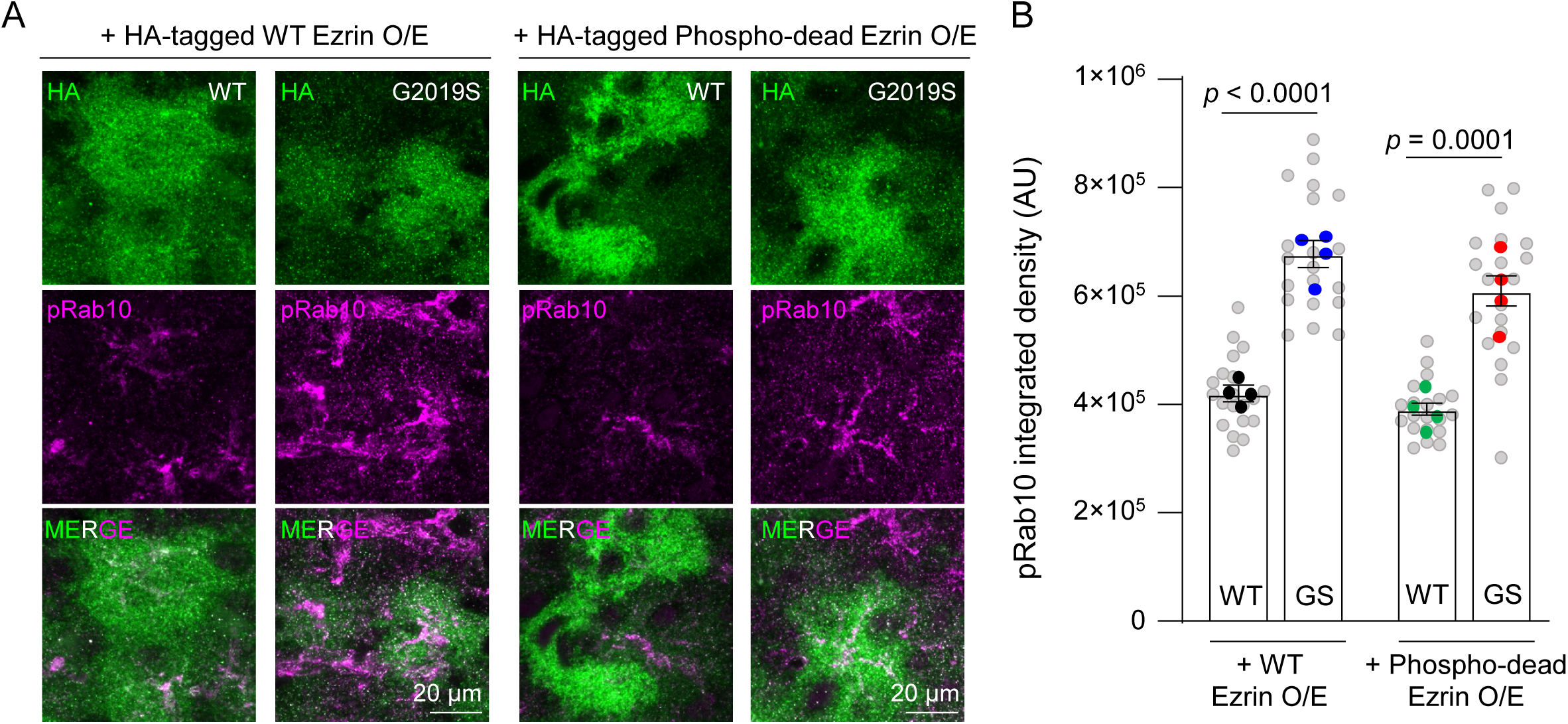
Overexpression of AAV-HA-tagged-WT Ezrin and AAV-HA-tagged-Phospho-dead Ezrin in WT and LRRK2 G2019S^ki/ki^ astrocytes. Related to Figure 4. **(A)** Representative images from the ventral ACC of WT and LRRK2 G2019S^ki/ki^ mice injected with AAV-HA-tagged WT Ezrin or Phospho-dead Ezrin, stained with HA and phospho-Rab10 antibodies at P84. Scale bar, 20 μm. **(B)** Quantification of phospho-Rab10 integrated density. n = 4 per group (2 males and 2 females). One-way ANOVA [F(3,12) = 38.34, *p* < 0.0001] with Bonferroni’s post hoc test revealed significant differences between groups, except WT + WT Ezrin O/E vs. WT + Phospho-dead Ezrin O/E (*p* > 0.9999) and LRRK2 G2019S^ki/ki^ groups (*p* = 0.3303).

**Figure S9 (related to Figure 4).**
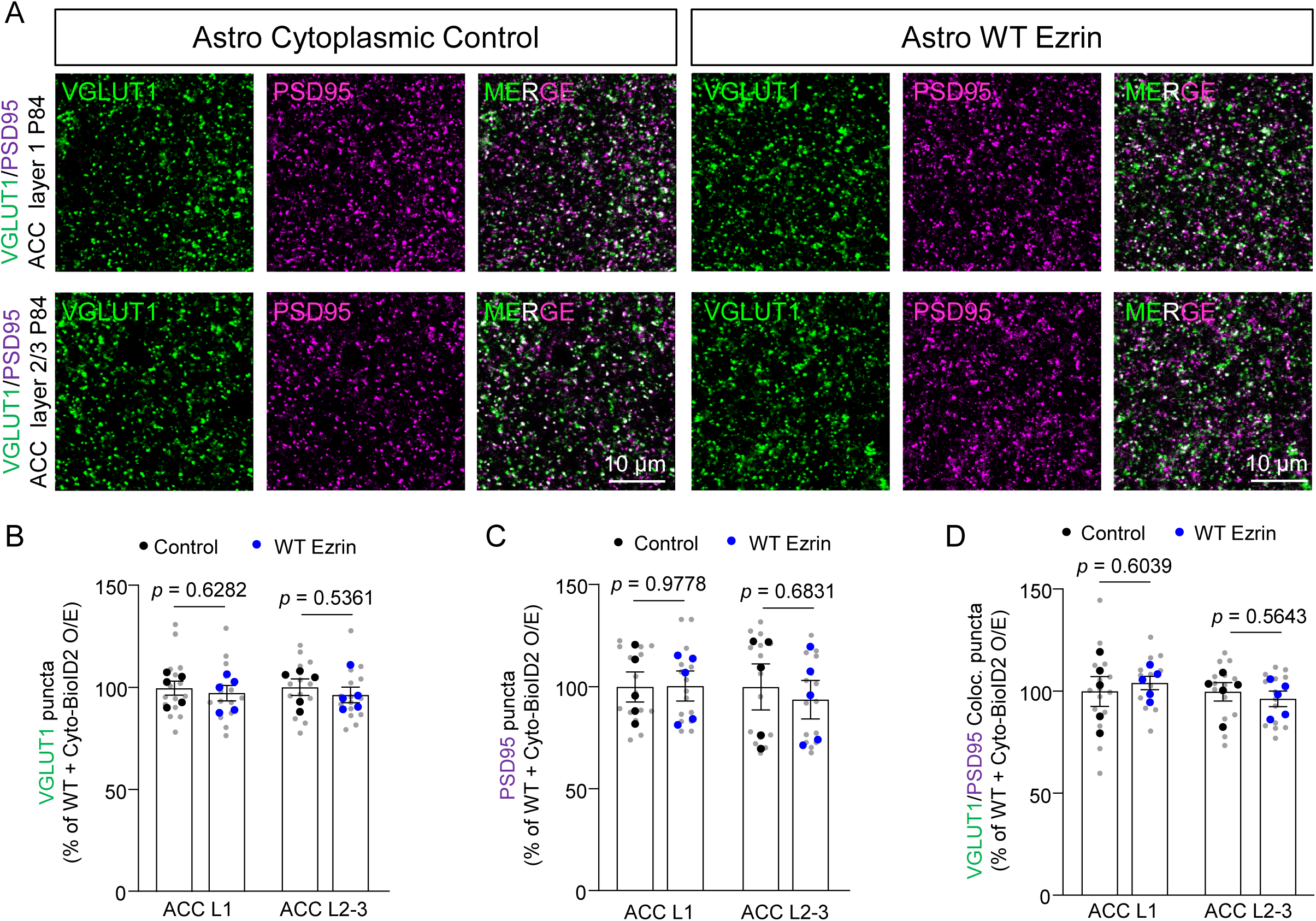
WT Ezrin overexpression does not change excitatory synapse numbers in the ACC of WT mice. Related to Figure 4. **(A)** Representative images from the ventral ACC of WT mice injected with AAV-HA-tagged BirA2 (cytoplasmic control) or WT Ezrin that were stained with VGLUT1 and PSD95 antibodies at P84. Scale bar, 10 μm. **(B)** Quantification of VGLUT1 puncta, normalized using the means of BirA2 values in the ventral ACC. n = 5 mice per group. For ACC L1, nested t-test, unpaired two-tailed t-test. t (8) = 0.5035, *p* = 0.6282. For ACC L2/3, nested t-test, unpaired two-tailed t-test. t (8) = 0.6465, *p* = 0.5361. **(C)** Quantification of PSD95 puncta, normalized using the means of BirA2 values in the ventral ACC. n = 5 mice per group. For ACC L1, nested t-test, unpaired two-tailed t-test. t (8) = 0.02872, *p* = 0.9778. For ACC L2/3, nested t-test, unpaired two-tailed t-test. t (8) = 0.4235, *p* = 0.6831. **(D)** Quantification of VGLUT1-PSD95 puncta, normalized using the means of BirA2 values in the ventral ACC. n = 5 mice per group. For ACC L1, nested t-test, unpaired two-tailed t-test. t (8) = 0.5400, *p* = 0.6039. For ACC L2/3, nested t-test, unpaired two-tailed t-test. t (8) = 0.6013, *p* = 0.5643.

**Figure S10 (related to Figure 4).**
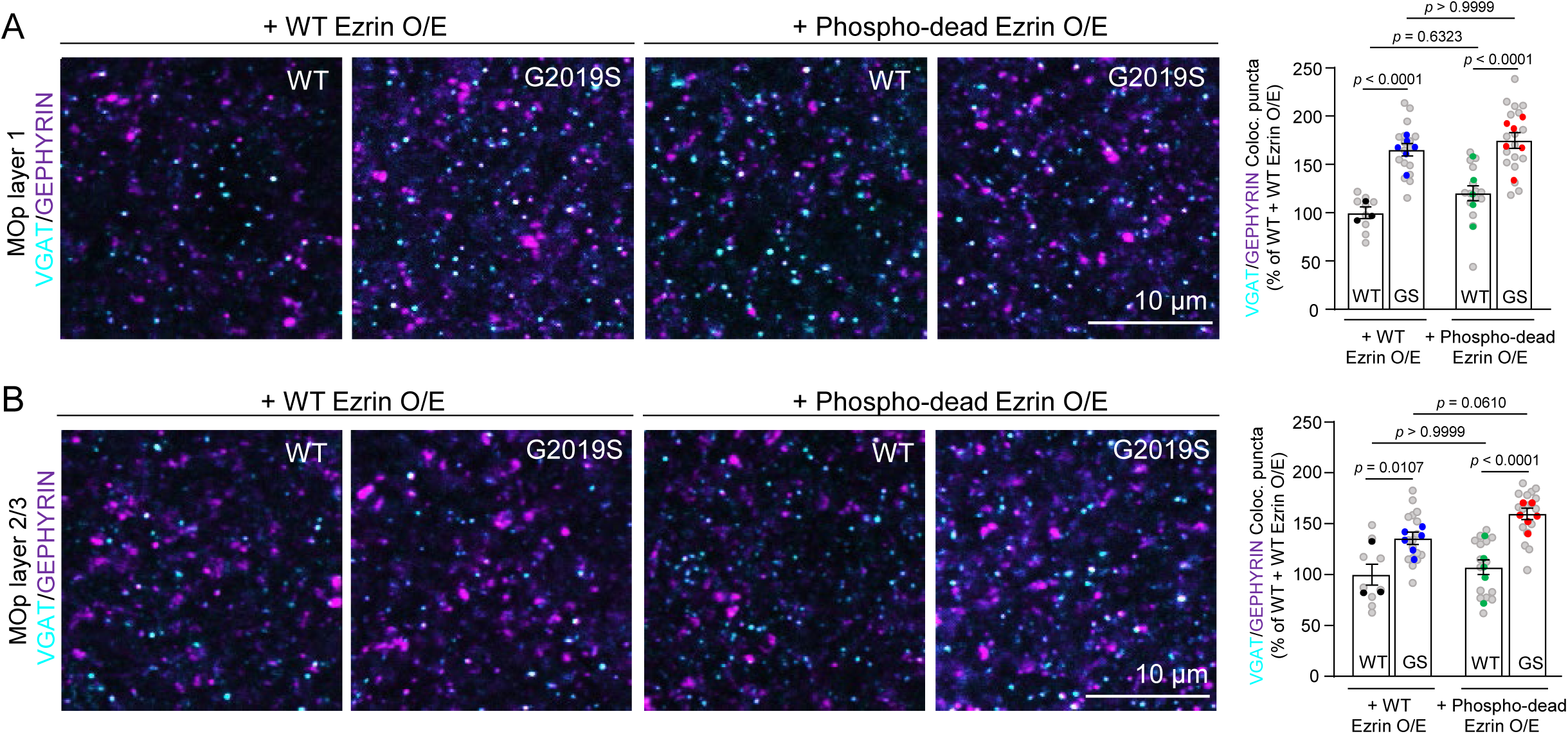
Overexpression of AAV-HA-tagged-WT Ezrin and AAV-HA-tagged-Phospho-dead Ezrin in WT and LRRK2 G2019S^ki/ki^ astrocytes. Related to Figure 4. **(A)** Representative images from the MOp L1 of the same groups stained with VGAT and GEPHYRIN antibodies. Scale bar, 10 μm. Quantification of VGAT-GEPHYRIN co-localized puncta, normalized to WT + WT Ezrin O/E. n = 3–6 per group. One-way ANOVA [F(3,55) = 19.7, *p* < 0.0001] with Bonferroni’s post hoc test revealed significant differences between WT + WT Ezrin O/E and LRRK2 G2019S^ki/ki^ + WT Ezrin O/E, and between WT + Phospho-dead Ezrin O/E and LRRK2 G2019S^ki/ki^ + Phospho-dead Ezrin O/E (*p* < 0.0001). No differences were found between WT or LRRK2 G2019S^ki/ki^ groups within Ezrin conditions (*p* > 0.05). **(B)** Representative images from the MOp L2-3 of WT and LRRK2 G2019S^ki/ki^ mice injected with AAV-HA-tagged WT Ezrin or Phospho-dead Ezrin, stained with VGAT and GEPHYRIN antibodies at P84. Scale bar, 10 μm. Quantification of VGAT-GEPHYRIN co-localized puncta normalized to WT + WT Ezrin O/E. n = 3–6 per group. One-way ANOVA [F(3,54) = 15.10, *p* < 0.0001] with Bonferroni’s post hoc test showed significant differences between WT + WT Ezrin O/E and LRRK2 G2019S^ki/ki^ + WT Ezrin O/E (*p* = 0.0107) and between WT + Phospho-dead Ezrin O/E and LRRK2 G2019S^ki/ki^ + Phospho-dead Ezrin O/E (*p* < 0.0001). No differences were observed between WT groups (*p* > 0.9999) or between LRRK2 G2019S^ki/ki^ groups (*p* = 0.0610).

**Figure S11 (related to Figure 5).**
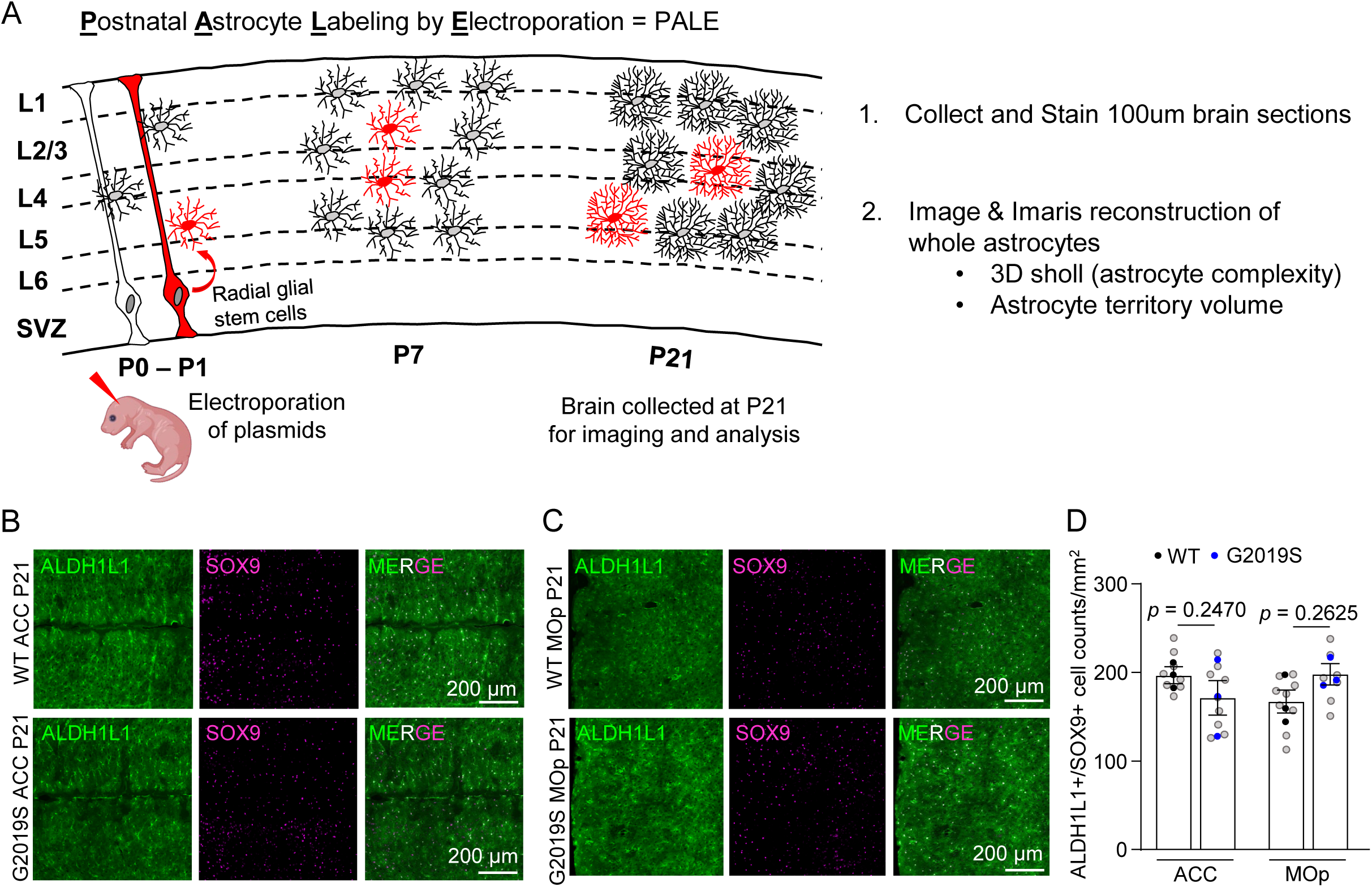
LRRK2 G2019S does not change ALDH1L1+/SOX9+ cell numbers in the ACC and MOp. Related to Figure 5. **(A)** Overview of postnatal astrocyte labeling by electroporation (PALE) with PiggyBac plasmids. **(B-C)** Representative confocal images of SOX9 (purple) in the ACC and MOp of WT or LRRK2 G2019S^ki/ki^ Aldh1l1-eGFP mice at P21. Scale bar, 200 μm. **(D)** Quantification of ALDH1L1+/SOX9+ cell numbers in (B-C), n = 3 (WT, 2 males and 1 female), 3 (LRRK2 G2019S^ki/ki^, 2 males and 1 female) mice, For ALDH1L1+/SOX9+ cell counting in the ACC, nested t-test, unpaired Two-tailed t-test. t (4) = 0.9619, *p* = 0.3906. For ALDH1L1+/SOX9+ cell counting in the MOp, nested t-test, unpaired two-tailed t-test. t (4) = 1.684, *p* = 0.1675. Grey dots are the data acquired from each image. Black dots are the averaged data acquired from each WT mouse. Blue dots are the averaged data acquired from each LRRK2 G2019S^ki/ki^ mouse.

**Figure S12 (related to Figure 5).**
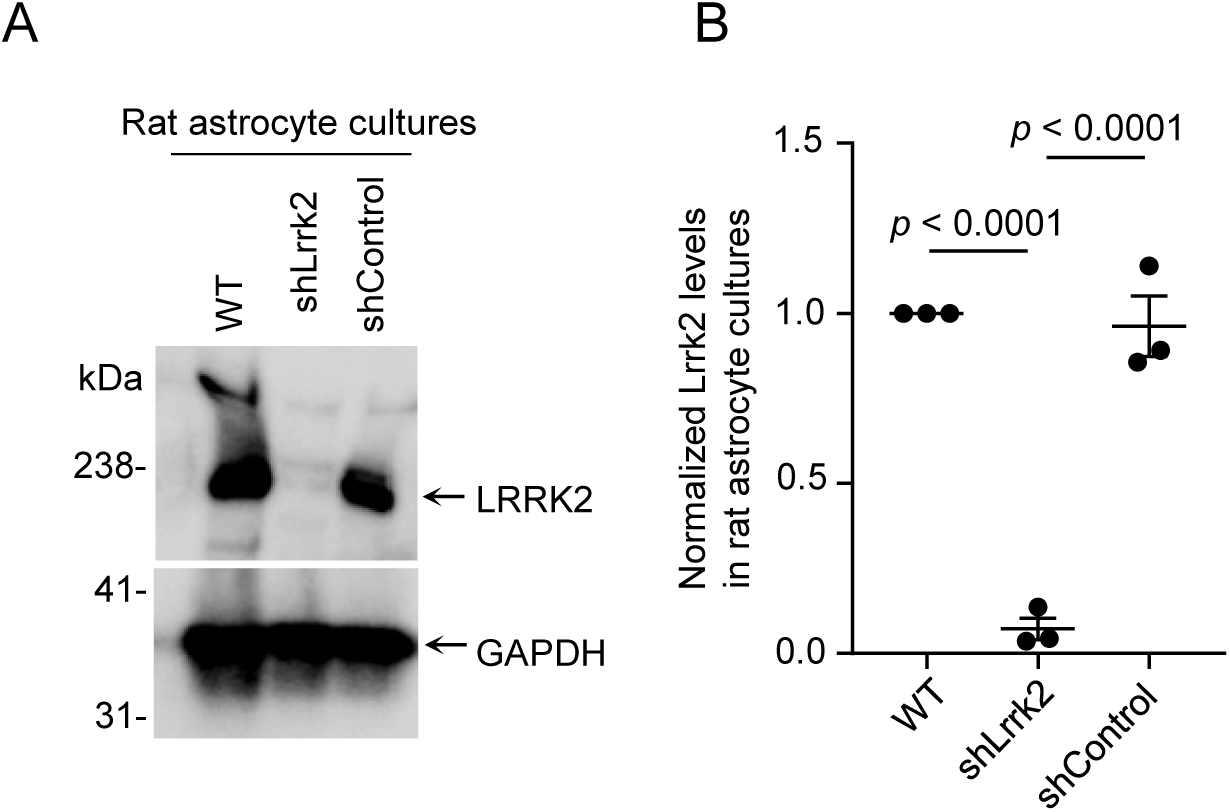
Expression of LRRK2 in astrocytes. Related to Figure 5. **(A)** Cultured WT, shControl, and shLrrk2 transfected rat cortical astrocytes were lysed, and cytoplasmic proteins were subjected to Western blotting using LRRK2 and GAPDH antibodies. Results are representative of 3 independent experiments. **(B)** Densitometric analysis of LRRK2 levels in (E). Signals corresponding to LRRK2 were first normalized to that for β-actin. Relative LRRK2 levels were then normalized to the LRRK2 signals in WT rat cortical astrocytes. Statistical significance was determined by One-way ANOVA [F (2, 6) = 92.25, *p* < 0.0001], Bonferroni multiple comparisons revealed a significant difference between WT and shLrrk2 (*p* < 0.0001, 95% C.I. = [0.6743, 1.183]) and between shLrrk2 and shControl (*p* < 0.0001, 95%C.I. = [-1.145, - 0.6363]) and no differences between WT and shControl (*p* > 0.9999, 95%C.I. = [-0.2165, 0.2923]), alpha = 0.05.

**Figure S13 (related to Figure 6).**
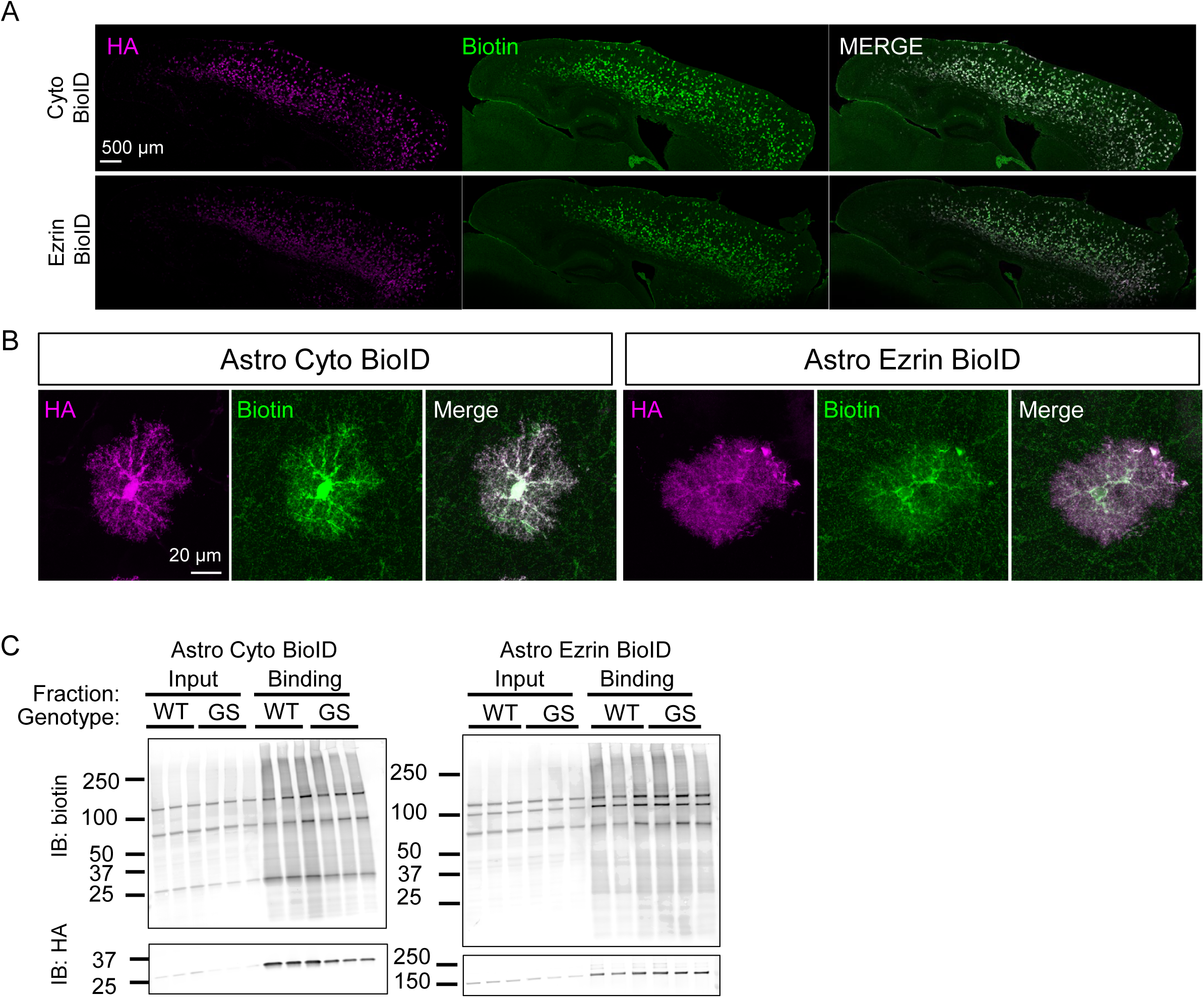
BioID constructs expression and biotinylation *in vivo* and *in vitro*. Related to Figure 6. **(A-B)** Representative images of *in vivo* expression in the cortex of different BioID constructs labeled with HA and the biotinylating activity labeled with streptavidin. Merged images show the colocalization of HA and biotin signals in astrocytes. Scale bar, 500 μm (A). Scale bar, 20 μm (B). **(C)** Western blot analysis of BioID constructs expression (HA) and biotinylation activity (Streptavidin) *in vitro* in cortical lysates and subsequent immunoprecipitation.

**Figure S14 (related to Figure 6).**
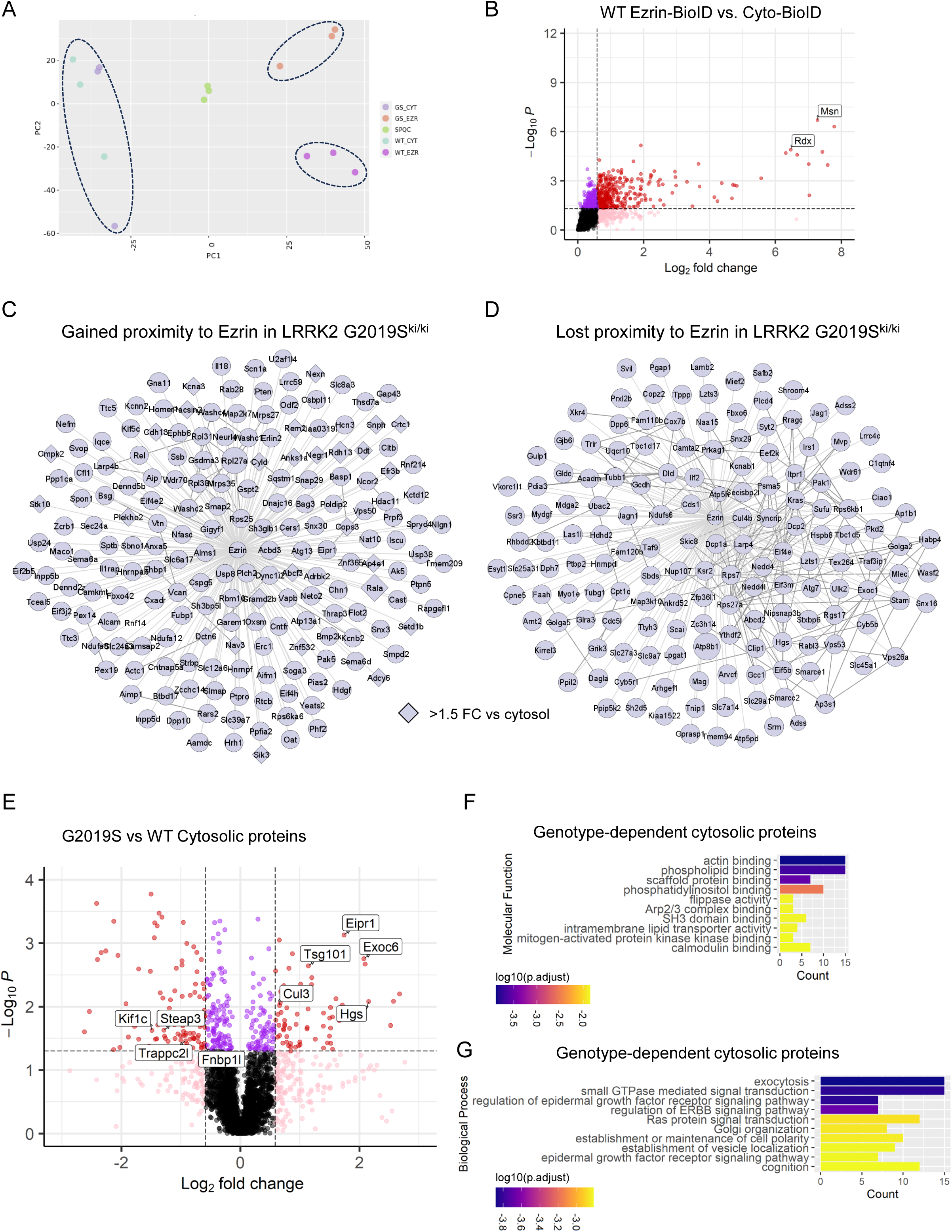
LRRK2 G2019S changes the composition of cytoplasmic proteins in astrocytes *in vivo*. Related to Figure 6. **(A)** Principle component analyses (PCA) of Cyto-BioID (control) and Ezrin-BioID (bait) samples. **(B)** The plot shows that Radixin and Moesin, known as Ezrin interacting proteins, are significantly enriched in the Ezrin-BioID sample compared to Cyto-BioID. **(C-D)** Interaction networks depict Ezrin interactions detected by BioID and known protein-protein interactions identified in the publicly available stringDB database for proteins with gained (C) or lost (D) proximity to Ezrin in LRRK2 G2019S^ki/ki^ astrocytes compared to WT astrocytes. **(E)** Volcano plot showing the differential abundance of cytosolic proteins in WT and LRRK2 G2019S^ki/ki^ cortices. **(F-G)** Bars show the top 10 most significant Gene Ontology (GO) terms, ordered by highest gene count and lowest adjusted p-value, for the proteins differentially detected by Astro-cyto-BioID in WT compared to LRRK2 G2019S^ki/ki^ (F) Molecular function (G) Biological Process.

**Figure S15 (related to Figure 7).**
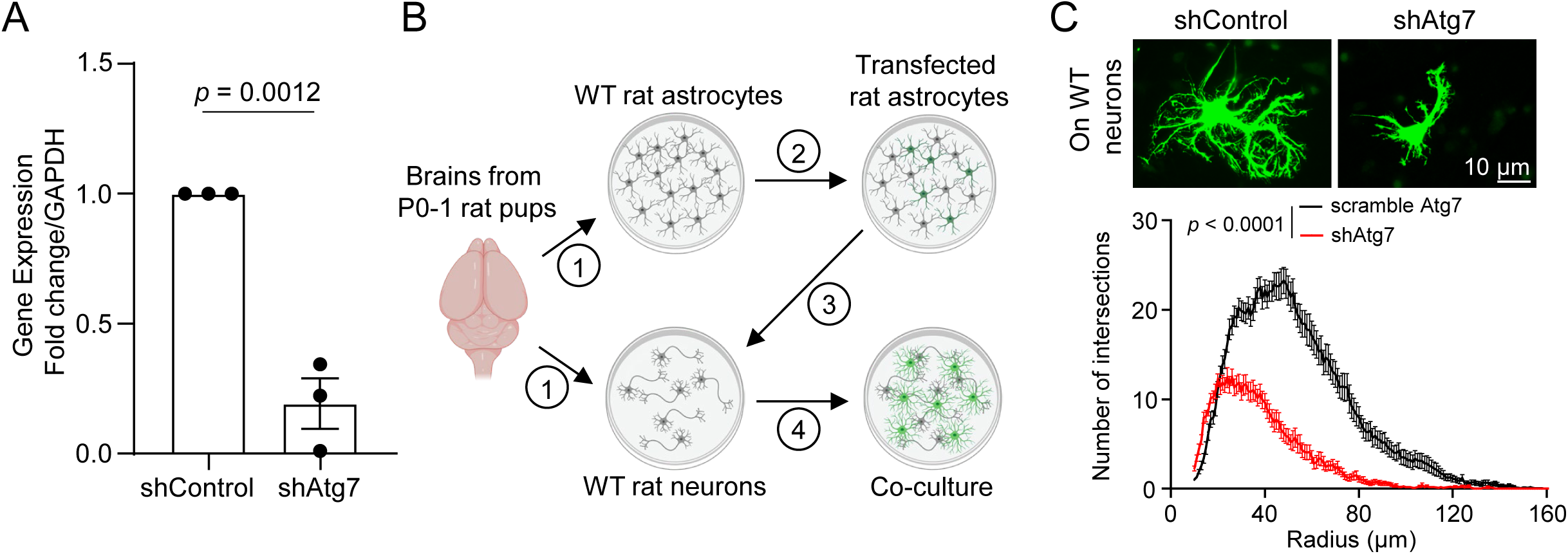
Astrocytic Atg7 is essential astrocyte morphological complexity *in vitro*. Related to Figure 7. **(A)** Comparable expression of Atg7 mRNA transcripts by RT-PCR in N2A cells transfected with scrambled shRNA (shControl-GFP) or shRNA targeting Atg7 (shAtg7-GFP). n = 3 independent cultures. unpaired Two-tailed t-test. t (4) = 8.258, *p* = 0.0012. **(B)** Schematic of astrocyte-neuron co-culture assay. **(C)** (Upper) Rat cortical astrocytes transfected with scrambled shRNA (shControl-GFP) or shRNA targeting Atg7 (shAtg7-GFP) co-cultured with wild-type cortical neurons. Scale bar, 10 μm. (Lower) Quantification of astrocyte branching complexity. n = 30 (shControl-GFP), 30 (shAtg7-GFP) astrocytes co-cultured with WT cortical neurons compiled from two independent experiments. We fitted a linear mixed model with a number of intersections as the outcome variable, condition as a predictor, and a number of cells and cell radius entered as random effects. Within this model, shAtg7 (beta = 2.91, t(58) = 7.14) led to a significant decrease in the number of intersections compared to shControl astrocytes (beta = 7.89, t(58) = 19.35). Tukey multiple comparisons test revealed a significant difference between shControl and shAtg7 astrocytes co-cultured on WT neurons (*p* < 0.0001).

